# Thresholds for post-rebound SHIV control after CCR5 gene-edited autologous hematopoietic cell transplantation

**DOI:** 10.1101/629717

**Authors:** E. Fabian Cardozo-Ojeda, Elizabeth R. Duke, Christopher W. Peterson, Daniel B. Reeves, Bryan T. Mayer, Hans-Peter Kiem, Joshua T. Schiffer

**Author notes:** To whom correspondence should be addressed. Address: 1100 Fairview Ave. N. MS E4-100, Seattle, WA, 98119.

## Abstract

Autologous, CCR5 gene-edited hematopoietic stem and progenitor cell (HSPC) transplantation is a promising strategy for achieving HIV remission. However, only a fraction of HSPCs can be edited *ex vivo* to provide protection against infection prior to autologous transplantation. The optimal transplantation conditions for achieving viral control in the absence of suppressive antiretroviral therapy (ART) are still unknown. We analyzed data from SHIV-1157ipd3N4-infected juvenile pig-tailed macaques that underwent autologous HSPC transplantation with and without CCR5 gene editing. We developed a mathematical model that recapitulates reconstitution of T cell subset counts and SHIV plasma viral loads in control and transplanted macaques. The model predicts that viral control can be obtained following ART treatment interruption (ATI) when: 1) levels of transplanted HSPCs are at least 10-fold higher than residual endogenous HSPCs after total body irradiation and 2) the fraction of protected HSPCs in the transplant achieves a threshold (73%-90%) sufficient to overcome transplantation-dependent loss of SHIV immunity. Under these conditions, if ATI is withheld until transplanted gene-modified cells engraft and reconstitute to a steady state, then spontaneous viral control is projected to occur immediately. Our results support strategies that 1) increase stem cell dose, 2) enhance potency of conditioning regimen, 3) elevate fraction of gene modified SHIV-resistant cells, 4) extend periods between HSPC transplantation and ATI with tracking of CD4^+^CCR5^-^ cell recovery and / or 5) augment anti-SHIV immunity to achieve sustained SHIV remission.

**One Sentence Summary:** Autologous transplantation of ΔCCR5 HSPCs may induce post-ATI SHIV control when the gene-edited cell dose is sufficient to overcome SHIV immunity loss.

## Introduction

The major obstacle to HIV-1 eradication is a latent reservoir of long-lived, infected cells (*1-3*). Cure strategies aim to eliminate all infected cells or permanently prevent viral reactivation from latency. The only known case of HIV cure (*4, 5*) and an additional recently-reported case of prolonged remission (*6*) resulted from allogeneic hematopoietic stem cell transplant with homozygous CCR5Δ32 donor cells (*4-6*). The success of this procedure is likely multifactorial— in part attributable to HIV resistance of the transplanted cells, the conditioning regimen that facilitates engraftment and eliminates infected cells, graft-versus-host effect against residual infected cells, and immunosuppressive therapies for graft-versus-host disease (*7-10*).

We are interested in recapitulating this method of cure with minimal toxicity. Specifically, we are investigating the use of autologous transplantation following *ex vivo* inactivation of the CCR5 gene with gene-editing (*11, 12*). This procedure is safe and feasible in pigtail macaques infected with simian-human immunodeficiency virus (SHIV) (*12-14*) and is currently being investigated in a Phase I clinical trial in suppressed, HIV-1-infected individuals (NCT02500849). Also, this approach is more broadly applicable because an allogeneic CCR5-negative donor is not needed. However, current data suggests that protocols do not achieve sufficient fractions of genetically modified HIV-resistant HSPCs. In contrast, in allogeneic transplant nearly 100% of circulating immune cells after engraftment consist of donor-derived CCR5Δ32 cells. This leads to a key question: what threshold percentage of CCR5-edited, autologous HSPCs is necessary for the cure/long term remission observed in the Berlin and London patients?

To answer this question, we developed a mathematical model that predicts the minimum threshold of gene-modified cells necessary for functional cure. First, we modeled the kinetics of CD4^+^CCR5^+^, CD4^+^ CCR5^-^, and CD8^+^ T cell reconstitution after autologous transplantation. Then we modeled SHIV kinetics during acute infection and rebound following ART treatment interruption to identify the degree of loss of anti-HIV cytolytic immunity following transplantation. Finally, we applied our models to predict the proportion of gene-modified cells necessary for stable virus remission, the dose of these cells relative to the intensity of the preparative conditioning regimen (total body irradiation, TBI), the timing of subsequent ATI, and the levels of SHIV-specific immunity required to maintain virus remission following ATI. Results from this three-part modeling approach support strategies that 1) increase stem cell dose, 2) enhance potency of conditioning regimen, 3) elevate fraction of gene modified SHIV-resistant cells, 4), extend periods between HSPC transplantation and ATI with tracking of CCR5-cell recovery and / or 5) augment anti-HIV immunity to achieve sustained HIV remission.

## Results

### Study design and mathematical modeling

We analyzed data from 22 juvenile pig-tailed macaques that were intravenously challenged with 9500 TCID50 SHIV1157ipd3N4 (SHIV-C) (**Fig. 1A**). After 6 months of infection, the macaques received combination ART that included tenofovir (PMPA), emtricitabine (FTC), and raltegravir (RAL). When on ART, 17/22 received total body irradiation (TBI) followed by the transplantation of autologous HSPCs with (n=12) or without (n=5) CCR5 gene editing (ΔCCR5 and WT groups, respectively). A control group (n=5) did not receive TBI or HSPC transplantation. 14 of the animals underwent ATI approximately one year after ART initiation. The remaining 8 animals were necropsied at an earlier time relative to the other animals’ ATI (see Methods for details).

**Figure 1.**
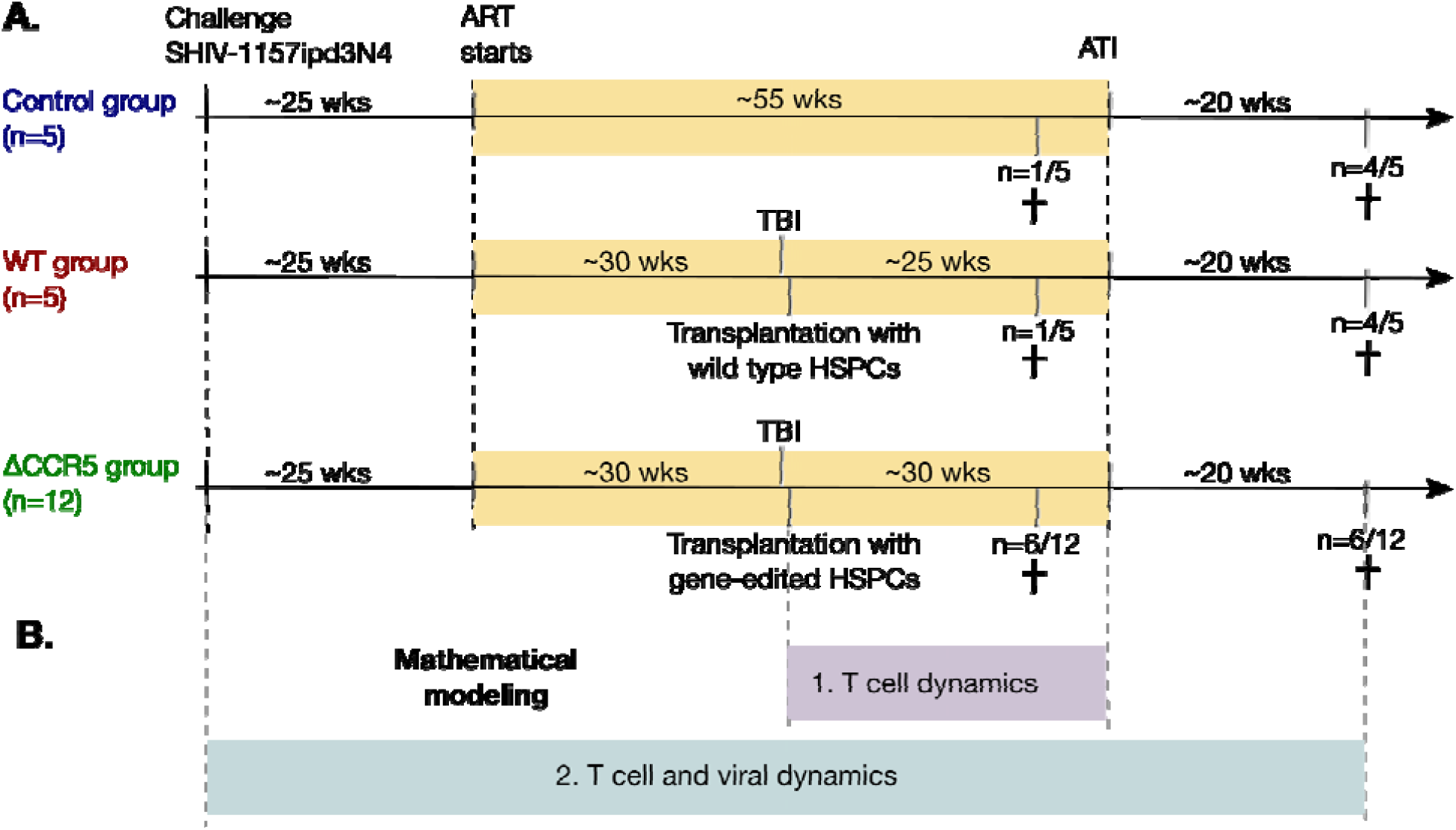
Study design and mathematical modeling. **A.** 22 pig-tailed macaques were infected with SHIV and suppressed with ART. Next, 17/22 underwent HSPC transplantation following myeloablative conditioning (TBI), including 12 animals that received CCR5-edited products and 5 that received non-edited products (ΔCCR5 and WT groups, respectively). A control group (n=5) did not receive TBI or HSPC transplantation. Fourteen animals underwent ATI approximately one year after ART initiation, while the remaining 8 animals were necropsied prior to ATI (see Methods for details). **B.** We first developed mathematical models for T cell dynamics and reconstitution following transplant and before ATI (purple), assuming that low viral loads on ART do not affect cell dynamics. After validation of that model, we introduced viral dynamics and fit those to the T cell and viral rebound dynamics from the animals pre- and post-ATI (blue).

To analyze the data and estimate thresholds for viral control under this approach we used ordinary differential equation models. We performed multi-stage modeling (**Fig. 1B**). First, we modeled the kinetics of CD4^+^ and CD8^+^ T cell subsets after autologous HSPC infusion following transplant and before ATI, assuming that ART suppression decouples SHIV-from cellular-dynamics. After validation of the first-stage model, we introduced a second-stage modeling to 1) explain virus and T cell kinetics during primary infection and analytical treatment interruption (ATI) and to 2) identify the degree of loss of anti-HIV cytolytic immunity due to the preparative conditioning. Then, we used the final validated model to project SHIV kinetics assuming different transplantation conditions.

### CD4^+^CCR5^+^ and CD8^+^ T cells recover more rapidly than CD4^+^CCR5^-^ T cells after HSPC transplantation

We analyzed the kinetics of peripheral blood CD4^+^CCR5^+^ and CD4^+^CCR5^-^ T cells, and total, T_naïve_, T_CM_, and T_EM_ CD8^+^ T cells in macaques after HSPC transplantation.

In untransplanted controls, levels of CD4^+^ and CD8^+^ T cells oscillated around a persistent set point (blue data-points in **Fig. 2**). Also, CD4^+^ CCR5^+^ T cell levels were ∼100 cells/μL and were uniformly lower than the CD4^+^CCR5^-^ T counts (each ∼1000 cells/μL) (**Fig. S1A**). Finally, total CD8^+^ T cell levels in the control group were ∼1400 cells/μl with a greater contribution from T_EM_ (73%) than T_N_+T_CM_ (27%) (based on median values, **Fig. S1B**).

**Figure 2.**
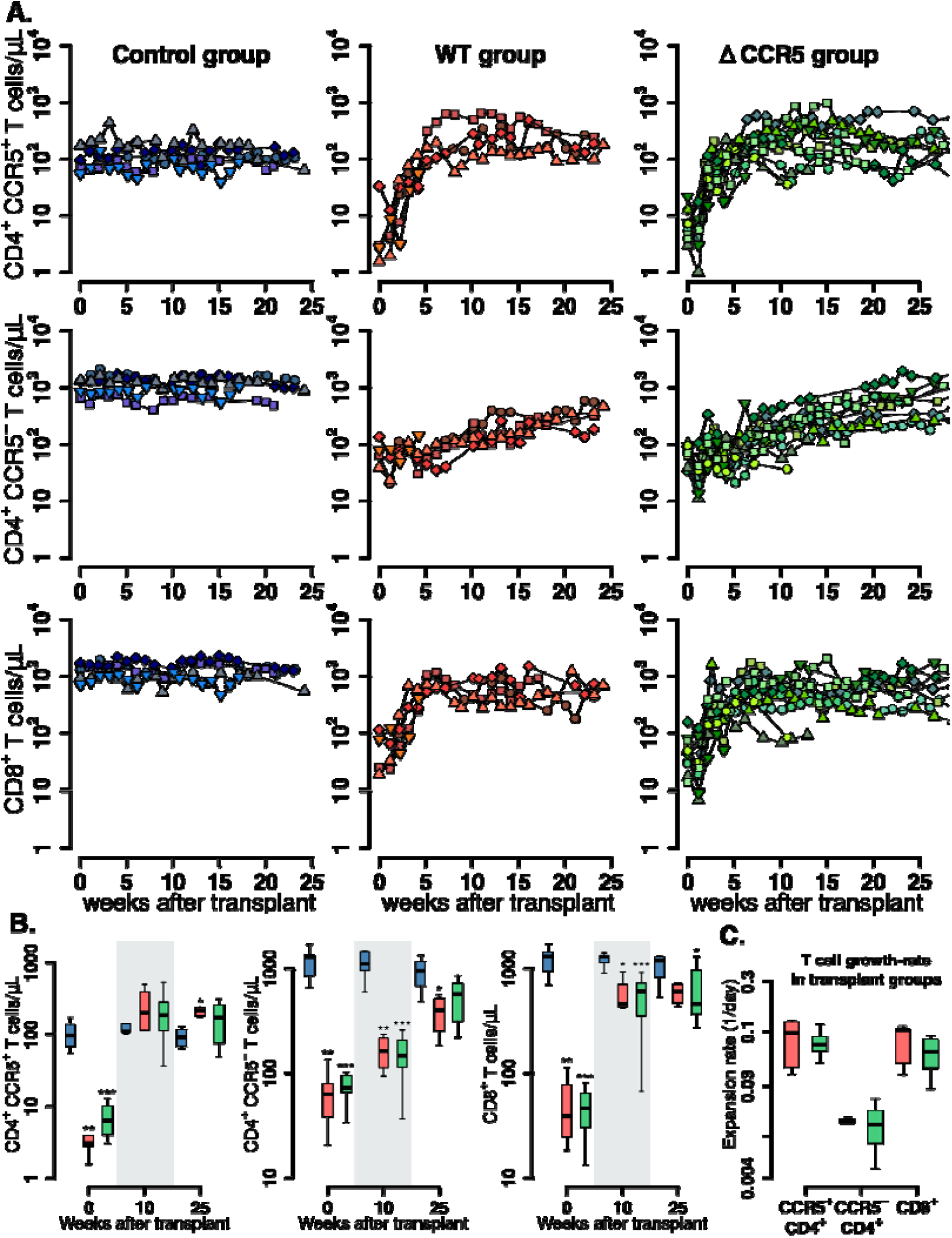
Post-transplantation, pre-ATI CD4^+^ and CD8^+^ T cell dynamics. **A.** Empirical data for peripheral CD4^+^ CCR5^+^ (top row), CD4^+^CCR5^-^ (middle row) and CD8^+^ T cell counts (bottom row) for control (blue), wild-type (red) and ΔCCR5 (green) transplantation groups. Each datapoint shape and color is a different animal sampled over time. **B.** Range of blood CD4^+^ and CD8^+^ T cell counts for weeks 0, 10 and 25 after transplantation (p-values calculated with Mann-Whitney test comparing control group with transplant groups. ^*^P<0.05, ^**^P<0.01 and ^***^P<0.001). **C.** Expansion-rate estimates of CD4^+^CCR5^+^, CD4^+^CCR5^-^, and CD8^+^ T cells (p-value calculated using a paired t-test). Colors for boxplots in **B & C** are matched to **A (**blue: control, red: wild-type-transplantation and green: ΔCCR5-transplantation groups).

In the transplant groups, post-TBI levels of CD4^+^CCR5^+^, CD4^+^CCR5^-^ and CD8^+^ T cells were significantly lower than in the control group but expanded at different rates during the following weeks (**Fig. 2A-C**). The levels of CD4^+^CCR5^+^ T cells started at 1-10 cells/μl and reconstituted to levels similar to the control group over 5-10 weeks (**Fig. 2A-B**). CD4^+^CCR5^-^ T cells remained at higher levels (∼100 cells/μL) than CD4^+^CCR5^+^ T cells after TBI but expanded more slowly and did not reach the values of the control group after 25 weeks (**Figs. 2A-B**). The CD4^+^CCR5^+^ T cell compartment expanded 8-fold more rapidly than the CD4^+^CCR5^-^ compartment (p=0.01, paired t-test, **Figs. 2C**). CD8^+^ T cells decreased to levels between 10 and 100 cells/μl after TBI but recovered to levels below the control group in 5 weeks (**Figs. 2A-B**); CD8^+^ T cells recovered as rapidly as the CD4^+^CCR5^+^ population (**Figs. 2C**).

Overall, these results show that after transplantation CD4^+^CCR5^+^ and CD8^+^ T cells recover faster than CD4^+^CCR5^-^ cells. This suggest that each cell subset may have different and/or complementary mechanisms that drive their expansion. To explore these mechanisms, we analyzed the data with a mechanistic mathematical model of cellular dynamics.

### Lymphopenia-induced proliferation drives early CD4^+^CCR5^+^ and CD8^+^ T cell reconstitution after HSPC transplantation

To identify the main drivers of T cell reconstitution after transplant, we developed a mathematical model that considered plausible mechanisms underlying reconstitution of distinct T cell subsets following autologous transplantation (**Fig. 3A**). We assumed that T cell reconstitution may have two main drivers: (1) lymphopenia-induced proliferation of mature cells that persists through myeloablative TBI (*15-19*), and (2) differentiation from naïve cells from progenitors in the thymus (from transplanted CD34^+^ HSPCs (*20, 21*) or residual endogenous CD34^+^ HSPCs that persist following TBI) and further differentiation to an activated effector state (*19, 22-26*). We also assumed the infused product dose *D* contains a fraction *f*_*p*_ of these transplanted, gene-edited HSPCs that do not express CCR5 (See **Table S1** for individual values of *D* and *f*_*p*_). Thus, in our model ΔCCR5-gene-modified CD4^+^ T cells differentiating from these modified HSPCs are subset of the total CD4^+^CCR5^-^ cell compartment (**Fig. 3A**), and therefore they have the same reconstitution kinetics.

**Figure 3.**
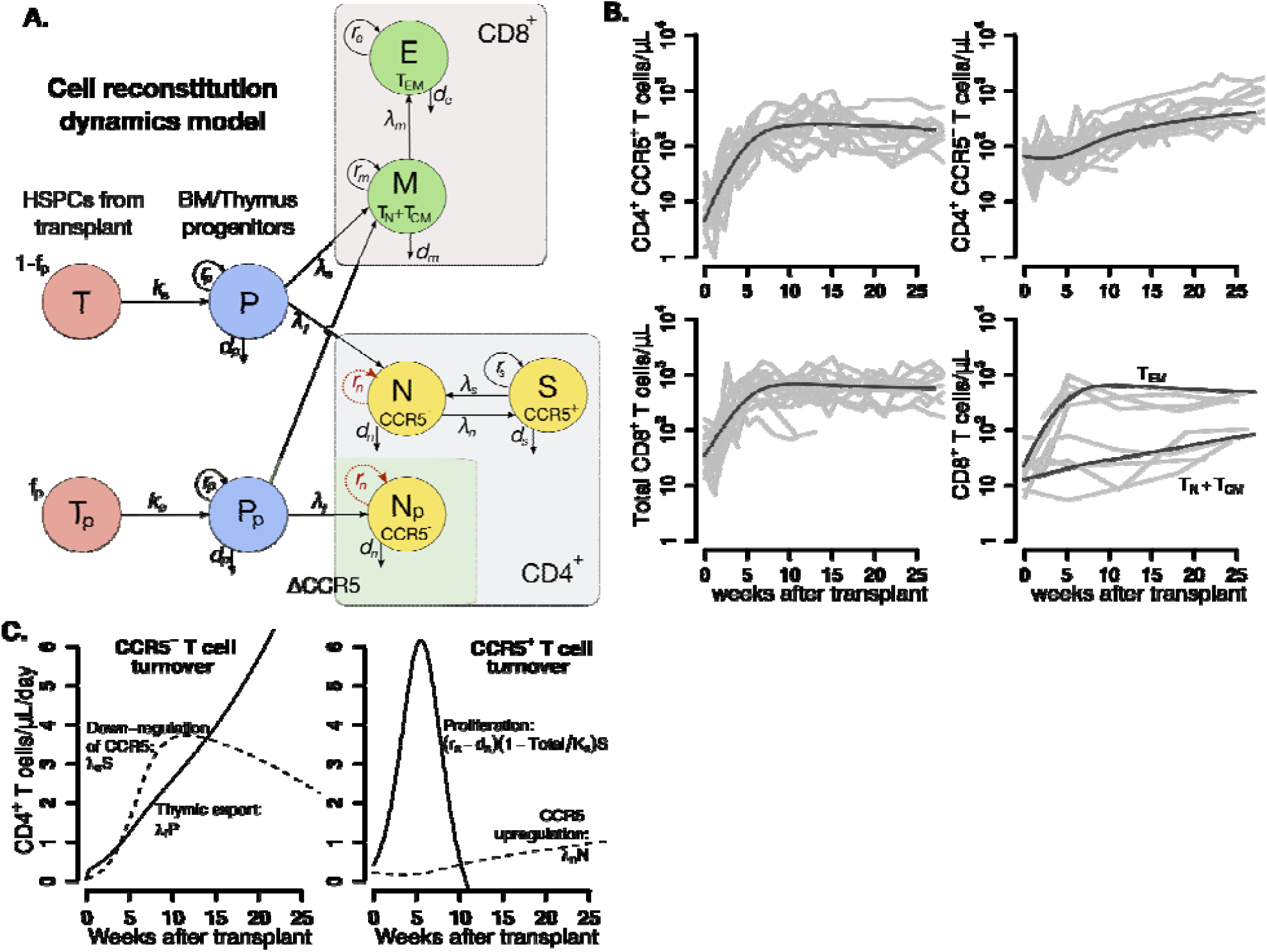
Mathematical model of T cell reconstitution after HSPC transplantation. **A.** Schematics of the model. Each circle represents a cell compartment: *T* represents the HSPCs from the transplant; *P*, the progenitor cells in bone marrow (BM) and thymus; *S* and *N*, CD4^+^CCR5^+^ and CD4^+^CCR5^-^ T cells, respectively; *T*_*p*_, the protected (ΔCCR5), gene-modified cells from transplant; *P*_*p*_, protected (ΔCCR5) progenitor cells in bone marrow/thymus; *N*_*p*_ the protected (ΔCCR5) CD4^+^ T cells; *M* the CD8^+^ T cells with naïve and central memory phenotype and *E* CD8^+^ T cells with effector memory phenotype. The initial fraction of protected cells is represented by the parameter *f*_*p*_. Gray panels represent mature blood CD4^+^ and CD8^+^ T cells, and green panel all ΔCCR5 cells in the model. Red, dashed arrows represent discarded terms after model selection and validation (see text for details). **B.** Model predictions using the maximum likelihood estimation of the population parameters (solid black lines) for all blood T cell subsets before ATI for all animals in the transplant groups. Each gray line is one animal. **C.** Model predictions of the total concentration of CD4^+^CCR5^-^ T cells generated by CCR5 downregulation (dashed line) or thymic export (solid line), and of the total concentration of CD4^+^CCR5^+^ T cells generated by proliferation (solid line) or by up-regulation of CCR5 (dashed line) over time using the maximum likelihood estimation of the population parameters.

We built 12 versions of the model by assuming that one or multiple mechanisms are absent, or by assuming certain mechanisms have equivalent or differing kinetics (**Table S2**). Using model selection theory, we identified the most parsimonious model that reproduced the data (schematic in **Fig. 3A** without red-dashed lines). The best model predictions for each cell subset using maximum likelihood estimates of the population parameters (**Table S3**) are presented in **Fig. 3B**. Individual fits are visualized in **Figs. S2-S4** and parameter estimates are collected in **Table S4**.

Model selection illuminated several biological phenomena: 1) CD4^+^CCR5^+^ T cell reconstitution after transplant is determined by cell proliferation and to a minor degree by upregulation of CCR5 (**Fig. 3C**); 2) CD4^+^CCR5^-^ T cell expansion is driven primarily by new naïve cells from the thymus and to a lesser extent by CCR5 downregulation (**Fig. 3C**); and 3) thymic export is not significantly different for CD4^+^ or CD8^+^ T cells (**Table S2**).

This first-stage modeling also makes testable biological predictions. First, the estimated CD4^+^CCR5^+^ T cell proliferation rate (∼0.1/day) far exceeds the estimated CCR5 upregulation (∼0.004/day) and thymic export rates (∼0.002/day). Therefore, one month after transplantation, the total concentration of CD4^+^CCR5^+^ T new cells generated by proliferation is predicted to be 40-fold higher than the concentration generated by up-regulation of CCR5 (**Fig. 3C**). Second, the CD8^+^ T_EM_ cells comprise the majority of the total CD8^+^ T cell compartment (**Fig. 3B**) with a proliferation rate up to 10-fold higher than the CD8^+^ T_CM_ cell differentiation rate (**Fig. S5**). In this way, CD8^+^ T cells follow a similar pattern to CD4^+^CCR5^+^ T cells (**Fig. 3B**).

In summary, following autologous HSPC transplant: 1) thymic export and downregulation of CCR5 drive a modest expansion of CD4^+^CCR5^-^ T cells, whereas 2) rapid lymphopenia-induced proliferation after TBI is the main driver for CD4^+^CCR5^+^ and CD8^+^ T cell expansion, which are derived from both the transplanted HSPC product and residual endogenous cells that persisted through the myeloablative conditioning regimen.

### Reduced CD4^+^CCR5^+^ T cell counts correlate with post-ATI plasma viral rebound only in animals that underwent HSPC transplantation without CCR5 editing

We compared plasma viral load rebound kinetics (*13, 27*) to CD4^+^CCR5^+^ and CCR5^-^ T cell subset dynamics after ART treatment interruption (ATI). **Fig. 4** presents the plasma viral loads and the blood CD4^+^CCR5^+^ and CD4^+^CCR5^-^ T cell kinetics before and after ATI in transplanted and control macaques. Viral burden after ATI was higher in the wild-type (WT) transplant group compared to the control and ΔCCR5 groups. Median peak viral load was 10-fold higher (p=0.06 comparing WT and control groups, Mann-Whitney test. **Fig. 4A**). Median viral load measurements at time of necropsy were 2-log_10_ higher (p=0.06 comparing WT and control groups, Mann-Whitney test. See **Fig.4B**). CD4^+^CCR5^+^ T cell counts decreased after ATI in the WT group, reaching a significantly lower nadir (∼8-fold) than in the control animals (p=0.01 comparing WT and control groups, Mann-Whitney test. **Fig. 4C**). The two animals with the largest reduction of CD4^+^CCR5^+^ T cells had the highest viral set points in the WT group. A similar trend was not observed in the ΔCCR5 group. There was no difference between CD4^+^CCR5^-^ T cell nadirs in the control and transplant groups (**Fig. 4D**).

**Figure 4.**
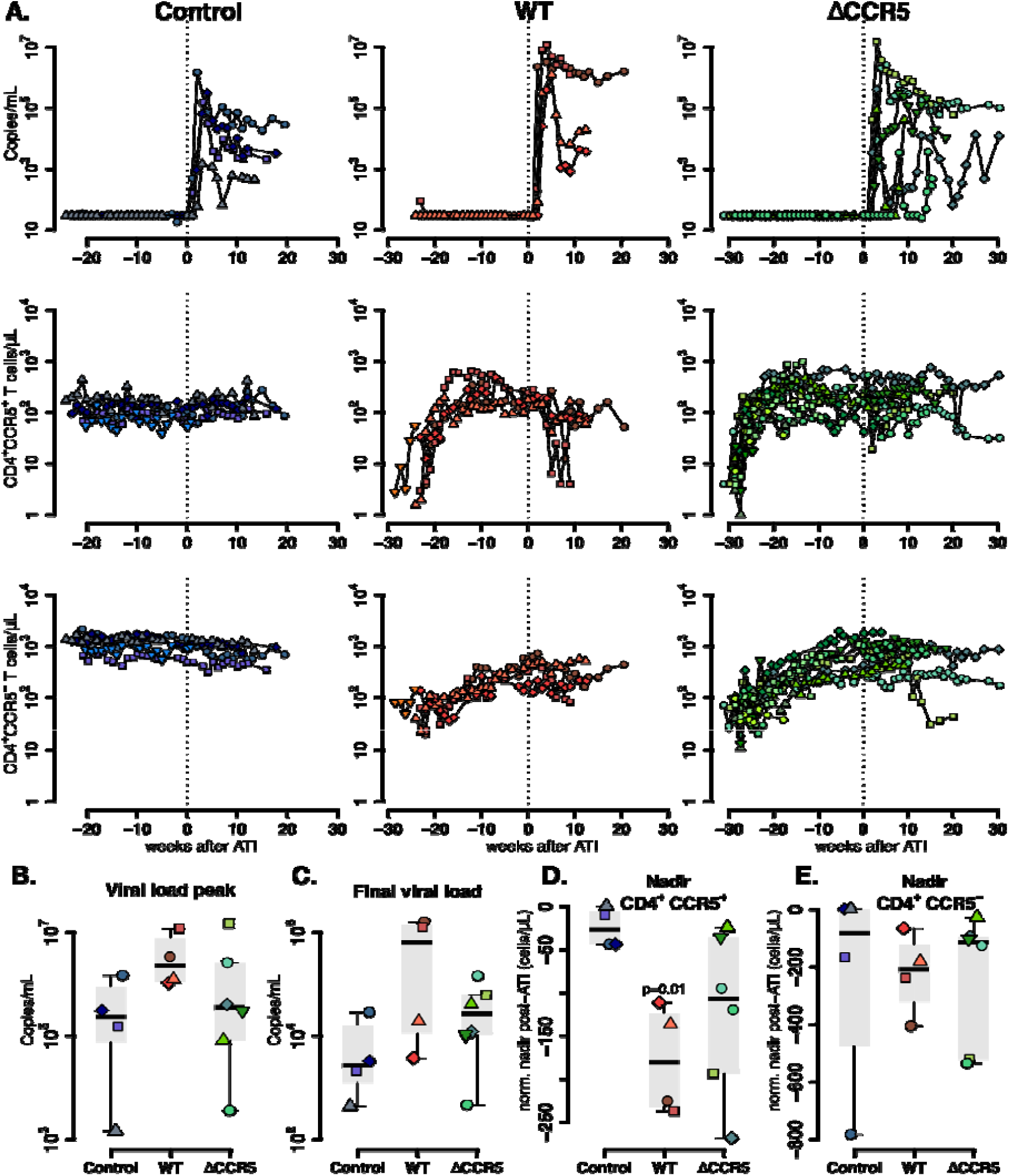
Plasma viral load and CD4^+^ T cell kinetics after ATI. **A.** Empirical data for viral load (top row) and peripheral T cell counts (middle and bottom rows) for control (blue), wild-type (red) and ΔCCR5 (green) transplantation groups. Each datapoint shape and color is a different animal sampled over time. **B-E**: Distributions of **B.** peak viral load post-ATI, **C.** viral load at endpoint necropsy, **D.** CD4^+^CCR5^+^ T-cell nadir post-ATI normalized relative to the CD4^+^CCR5^+^ concentration at ATI, **E.** CD4^+^CCR5^-^ T-cell nadir post-ATI normalized relative to the CD4^+^CCR5^-^ concentration at ATI. Controls were compared to transplant groups using Mann-Whitney test for all summary measures.

In the control and ΔCCR5 groups individual viral loads did not correlate with corresponding CD4^+^CCR5^+^ T cell counts post-ATI. However, in three out of the four animals in the wild-type transplant group, viral load observations post-ATI correlated negatively with their corresponding CD4^+^CCR5^+^ T cell counts (**Fig. S6**). To summarize, only animals that were transplanted with wild-type HSPCs had higher viral loads that correlated with reductions in CD4^+^CCR5^+^ T cells and more rapid disease progression after ATI. When CCR5-editing was included in the transplanted HSPC product, viral rebound and the degree of T cell depletion was more similar to the control group (no transplantation). We hypothesize that transplantation augments depletion of CD4^+^CCR5^+^ T cells following ATI, which can be rescued via ΔCCR5 edition of the HSPC product.

### A reduction in SHIV-specific immunity leads to higher viral rebound set points and CD4^+^CCR5^+^ T cell depletion following ATI in transplanted animals

We simultaneously analyzed the viral and T cell subset data using mechanistic mathematical models in order to understand how transplantation modifies plasma viral load and CD4^+^CCR5^+^ T cell kinetics during ATI. We extended our T cell reconstitution model to include SHIV infection (**Fig. 5A** and **Methods**) and used this second-stage model to analyze virus and T cell dynamics from SHIV-infection until ATI. Again following model selection theory based on AIC, we found the most parsimonious model to explain the data (**Fig. 5A, Table S5**). This model simultaneously recapitulates plasma viral load and the kinetics of CD4^+^ CCR5^+^ and CCR5^-^ T cells during primary infection and rebound after ATI in each animal as shown in **Fig. 5B** and **Figs. S7-S9** with corresponding estimated parameters in **Tables S6-S7**. In this model, SHIV-specific CD8^+^ effector cells reduce virus production rather than killing infected cells (*28-30*), possibly by secretion of HIV-antiviral factors (*31-33*)—not explicitly included in the model. The model also suggests that infection enhances upregulation of CD4^+^CCR5^-^ T cells. This upregulation replenishes CD4^+^CCR5^+^ T cells and transiently reduces the CD4^+^CCR5^-^ compartment after ATI (*34-36*). Finally, the data-fit required different SHIV-specific CD8^+^ T cell proliferation, saturation and death rates, i.e. parameters and *ω*_8,_ *I*_50_ and *d*_*h*_ from **eq. 3**, respectively between ATI and acute infection (see **Table S5**) (*27*).

**Figure 5.**
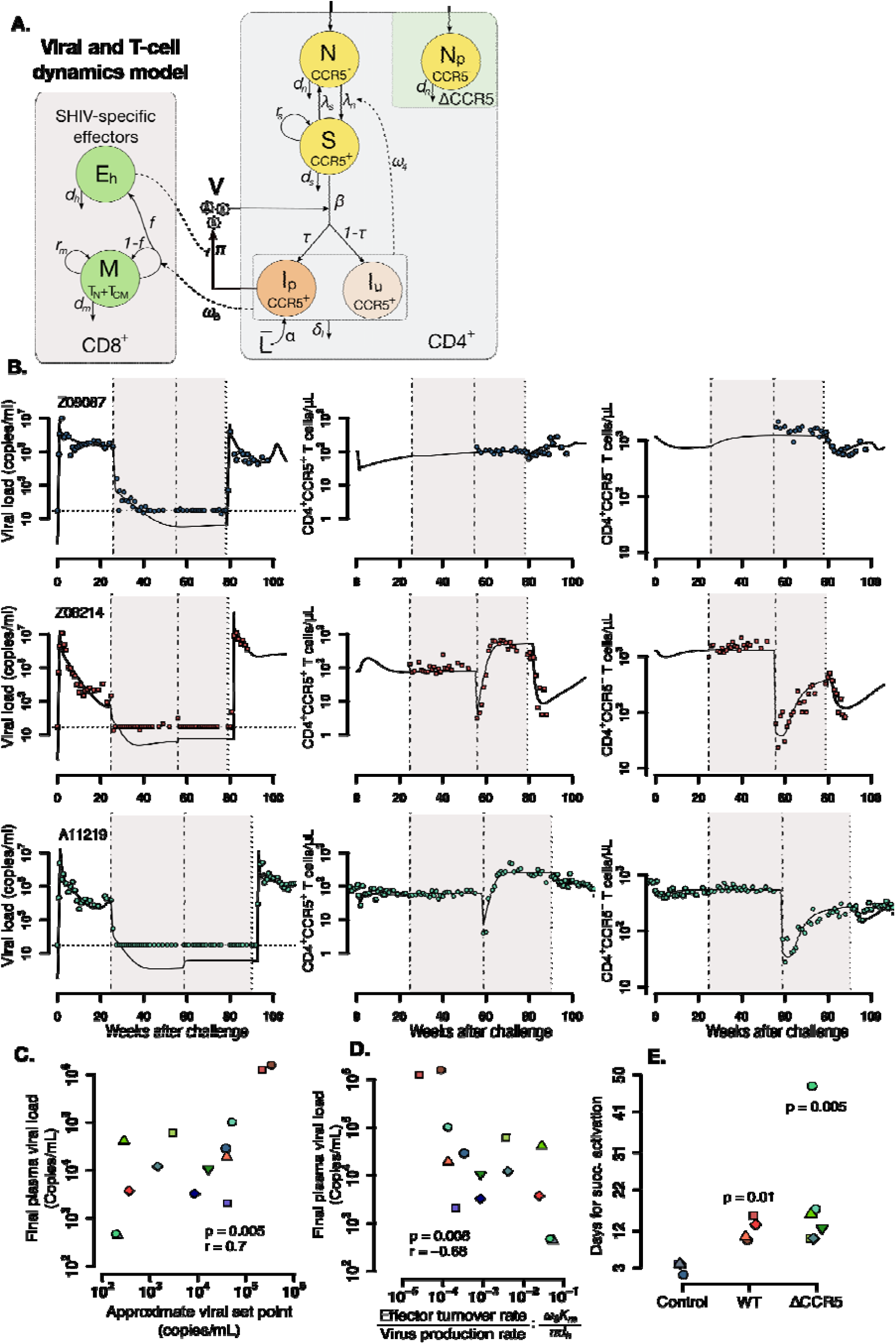
Mathematical model of virus and T cell dynamics following ATI. **A.** Model: Susceptible cells, *S*, are infected by the virus, *V*, at rate *β. I*_*p*_ represents the fraction *τ* of the infected cells that produce virus, and, *I*_*u*_, the other fraction that becomes unproductively infected. Total CD4^+^CCR5^+^ T cell count is given by the sum of *S, I*_*p*_ and *I*_*u*_. All infected cells die at rate Δ_*I*_. *I*_*P*_ cells arise from activation of latently infected cells at rate 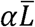 and produce at a rate *π*; virus that is cleared at rate *γ*. CD8^+^ *M* cells proliferate in the presence of infection with rate *ω*_*8*_ from which a fraction *f* become SHIV-specific CD8^+^ effector T cells, *E*_*h*_, that are removed at a rate *d*_*h*_. These effector cells reduce virus production (π) by 1/(1+θ*E*_*h*_). Non-susceptible CD4^+^ T cells that were not CCR5-edited upregulate CCR5 in the presence of infection and replenish the susceptible pool at rate *ω*_*4*_. Gray panels represent mature blood CD4^+^ and CD8^+^ T cells, and green panel ΔCCR5 cells in the model. **B.** Individual fits of the model (black lines) to SHIV RNA (left column), blood CD4^+^CCR5^+^ T cells (middle column) and CD4^+^CCR5^-^ T cells (right column) for one animal in the control (top row), wild type (middle row) and ΔCCR5 groups (bottom row). Shaded areas represent time during ART and dashed-point line, the time of transplantation. **C-D:** Scatterplots of final observed viral load vs. **C.** Approximate viral set points obtained by the model, and **D.** the ratio between the turnover rate of SHIV-specific CD8^+^ T cells and the virus production rate: 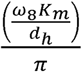 (p-values calculated using Pearson’s correlation test). **E.** Individual estimates of the time until SHIV successful activation after ATI (see **Methods**). Blue: control, red: wild type, and green: ΔCCR5 transplant group (p-values calculated by Mann-Whitney test).

We used our model to calculate an approximate viral load set point (**Fig. 5C** and **Methods**) after treatment interruption. This set point is driven by the ratio between the turnover rate of SHIV-specific CD8^+^ T cells and the virus production rate: 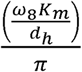 (see **Methods**). This ratio correlates negatively with the measured final viral load (**Fig. 5D**). In this sense, the highest viral set point in animals in the transplant groups might be due to the underlying low immune response to the virus. However, we found that no individual SHIV-specific immunity parameter was correlated with viral set point or was different among the three groups. Rather, the only estimated parameter that differed among groups was the time until successful SHIV reactivation (*t*_*sa*_, see **Methods**) which was significantly higher in the transplant groups (**Fig. 5E**). In conclusion, we developed a second-stage model that simultaneously recapitulates viral and T cell dynamics from SHIV-infected animals receiving autologous HSPC transplantation. This model implies that transplant reduces host T-cell immunity, resulting in higher viral loads after ATI in transplanted animals.

### Post-ATI viral control requires a large HSPC dose containing a high fraction of CCR5-edited cells

An important advantage of our model is the ability to calculate the conditions required for post-ATI viral control (viral load set point < 30 copies/ml) after CCR5-edited autologous transplant. To this end, we used our second-stage model to approximate an effective reproductive ratio *R*_*eff*_to describe the ability of the virus to sustain infection after ATI (see **Methods**):

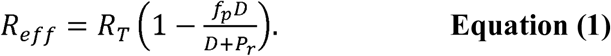

Here, *f*_*p*_ describes the fraction of protected HSPCs in the transplant product, *D* the dose or total number of infused HSPCs, and *P*_*r*_ the number of residual endogenous HSPCs after conditioning (variable *P*, **Fig. 3A**). *R*_*T*_ is the approximate number of new infections caused by one infected cell after T cell complete reconstitution post-conditioning as defined in **eq. 4** (see Methods) and is inversely related to the anti-SHIV immune response at the time of ATI. Post-ATI viral control depends on the number of HSPCs in the body immediately after transplant that are protected from SHIV infection, or 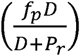.

To estimate the values of *f*_*p*_, *D*, and *P*_*r*_ needed for viral control, we first estimated *R*_*T*_ for each animal based on individual parameter estimates pertaining to anti-SHIV immunity. We then simulated the model for each animal using varying values of *f*_*p*_ from zero to one (0-100% CCR5-edited HSPCs), values of *D* from 10^6^ to 10^9^ HSPCs, and values of *P*_*r*_ from zero to 10^7^ HSPCs. As an illustration, **Fig. 6A** depicts projections of the model for transplanted animal A11219 for selected values of *f*_*p*_ when *D* = 10^7^ HSPCs/kg and *P*_*r*_ = 6 × 10^6^ HSPCs. When *f*_*p*_, *D* and *P*_*r*_, made *R*_*eff*_ ≥ 1, plasma viremia rebounded following ATI and virus was not controlled. When *R*_*eff*_ < 1, post-rebound control was observed, but only at weeks 40-60 post-ATI, following an initial decrease in viral loads beginning 30-40 weeks after ATI. In this case, post-rebound control occurred concomitantly with ΔCCR5 CD4^+^ T cell complete reconstitution relative to non-edited CCR5^+/-^ CD4^+^ T cells (**Fig 6B**). Lower values of *R*_*eff*_ resulted in earlier post-rebound control.

**Figure 6.**
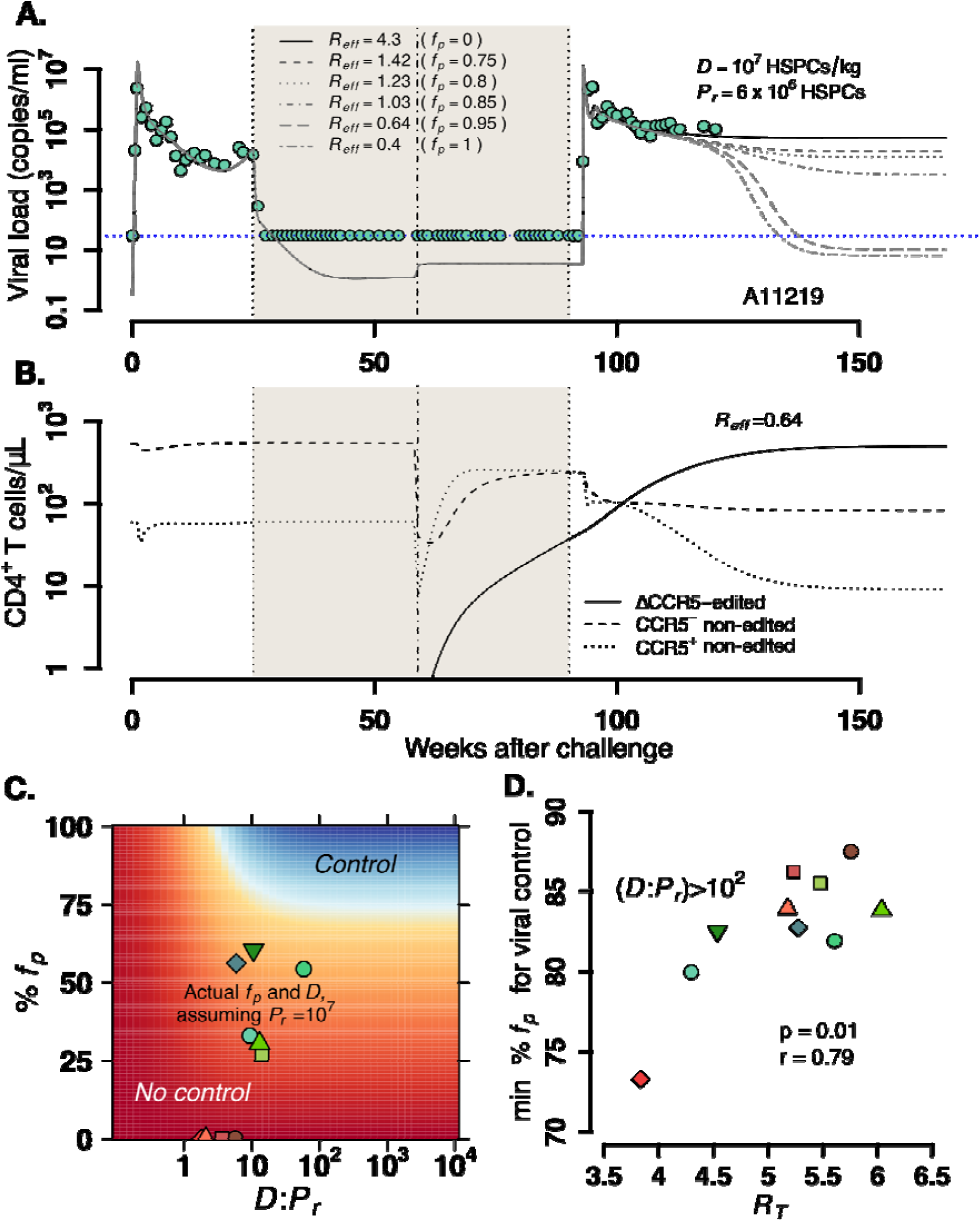
Model predictions of factors governing post-rebound viral control after CCR5 gene-edited HSPC transplant. **A.**Predictions for plasma viral loads post-ATI using the optimized mathematical model. Here, — and is the composite determinant of viral control. Parameter estimates for animal A11219 (**Table S7)** were used to compute the effective reproductive ratio *R*_*T*_. Higher values of *R*_*T*_ imply poorer anti-SHIV immunity (see **eq. 4** in Methods). We varied values of the fraction of HSPCs in transplant *f*_*p*_, the stem cell dose *D* as shown and fixed the remaining number of HSPCs after TBI before transplant *P*_*r*_ = 6 ×10^6^. *R*_*eff*_ <1 predicted spontaneous viral control 40-50 weeks after ATI. **B.** A simulation with *R*_*eff*_ = 0.64 demonstrates CCR5-edited CD4+ T cell recovery is concurrent with viral control. **C.** Model predictions of the fraction of protected HSPCs in the transplant *f*_*p*_ (y-axis) and the fraction of transplanted HSPCs with respect to the total infused plus remaining post-TBI HSPCs *D*: *P*_*r*_ (x-axis) required for spontaneous viral control. *R*_*T*_ =, 3.7 is from animal A11200 which has the lowest *f*_*p*_ (74%) and *D*: *P*_*r*_ (∼10) required for post-ATI viral control (heatmaps for other animals in **Fig. S11**). Blue color represents the parameter space with post-ATI viral control or *R*_*eff*_ <1. Yellow-to-red colors represent the parameter space with no control or *R*_*eff*_ >1. Data points represent the individual values of *f*_*p*_ and *D*: *P*_*r*_ from each transplanted animal in the study. **D.** Model predictions of the minimum fraction of protected HSPCs in the body *f*_*p*_ for viral control (y-axis) when 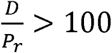 is assumed for each animal given their calculated values for *R*_*T*_ (x-axis). Each color is an animal, and A11200 is the red diamond with the lowest value of min *f*_*p*_. A lower *f*_*p*_ may lead to viral control if there is a more potent anti-SHIV immune response.

For all animals, post-treatment control occurred when values of *f*_*p*_, *D* and *P*_*r*_ made *R*_*eff*_ < 1(**Fig. S10**). Model predictions of animal A11200 demonstrate that regardless of the fraction of protected HSPCs in the transplant (*f*_*p*_), viral control is possible only when the ratio of HSPCs in the transplant to the residual endogenous HSPCs in the body post-TBI (*D* : *P*_*r*_) is above 10 (**Fig. 6C**). Moreover, if the ratio *D* : *P*_*r*_ is greater than 10, the minimum fraction of protected cells required is 73%. When the ratio *D*: *P*_*r*_ is higher than 100, we found that the minimum fraction of protected cells in the transplant *f*_*p*_ varied from 73% to 90% in all transplanted animals and was positively correlated with a weaker anti-SHIV immune response of the given animal defined by *R*_*T*_ (**Fig. 6D**). Similar results were obtained for all other animals (**Fig. S11**). This is consistent with **eq. 1** as *R*_*eff*_ ≈ *R*_*T*_ (1 − *f*_*p*_) when *D* ≫ *P*_*r*_. *R*_*T*_ varied from 3 and 6 across animals using individual parameter estimates in **Table S7**. The required levels for *f*_*p*_ are lower in the context of more intense anti-SHIV immunologic pressure. This result argues for strategies that 1) sustain anti-SHIV immunity (lower *R*_*T*_), 2) increase the stem cell dose relative to the residual endogenous stem cells (*D*: *P*_*r*_) after transplant—perhaps by enhancing potency of the conditioning regimen, and 3) increase the fraction of gene-modified, SHIV-resistant cells (*f*_*p*_).

Based on the observation the viral control occurred when CD4^+^ T cell subsets reached a steady state in the simulations (**Fig. 6A-B)**, we simulated the model again for animal A11219 under conditions that lead to viral control: *f*_*p*_ = 0.95, *D* = 10^8.5^ HSPCs and *P*_*r*_ = 10^7^ HSPC with ATI occurring at 3, 14, 26 or 38 weeks after transplantation. Indeed, time to post-ATI viral control (shaded areas in **Fig. 7A**) decreased as time to ATI was extended after transplant and as the difference between CD4^+^CCR5^-^ cell density at ATI and its expected set point decreased (shaded areas in **Fig. 7B**). In this case, ΔCCR5 CD4^+^ T cells comprised the majority of the CD4^+^CCR5^-^ T cell compartment (**Fig. 7B**). When including varying times of ATI from 0 to 60 weeks after transplantation, the model predicted the time between transplant and ATI required to avoid viral rebound is about 30 weeks for animal A11219 (**Fig. 7C**). This timeframe allowed a CD4^+^CCR5^-^ cell density at ATI exceeding 80% of the ultimate steady state value (**Fig. 7D**).

**Figure 7.**
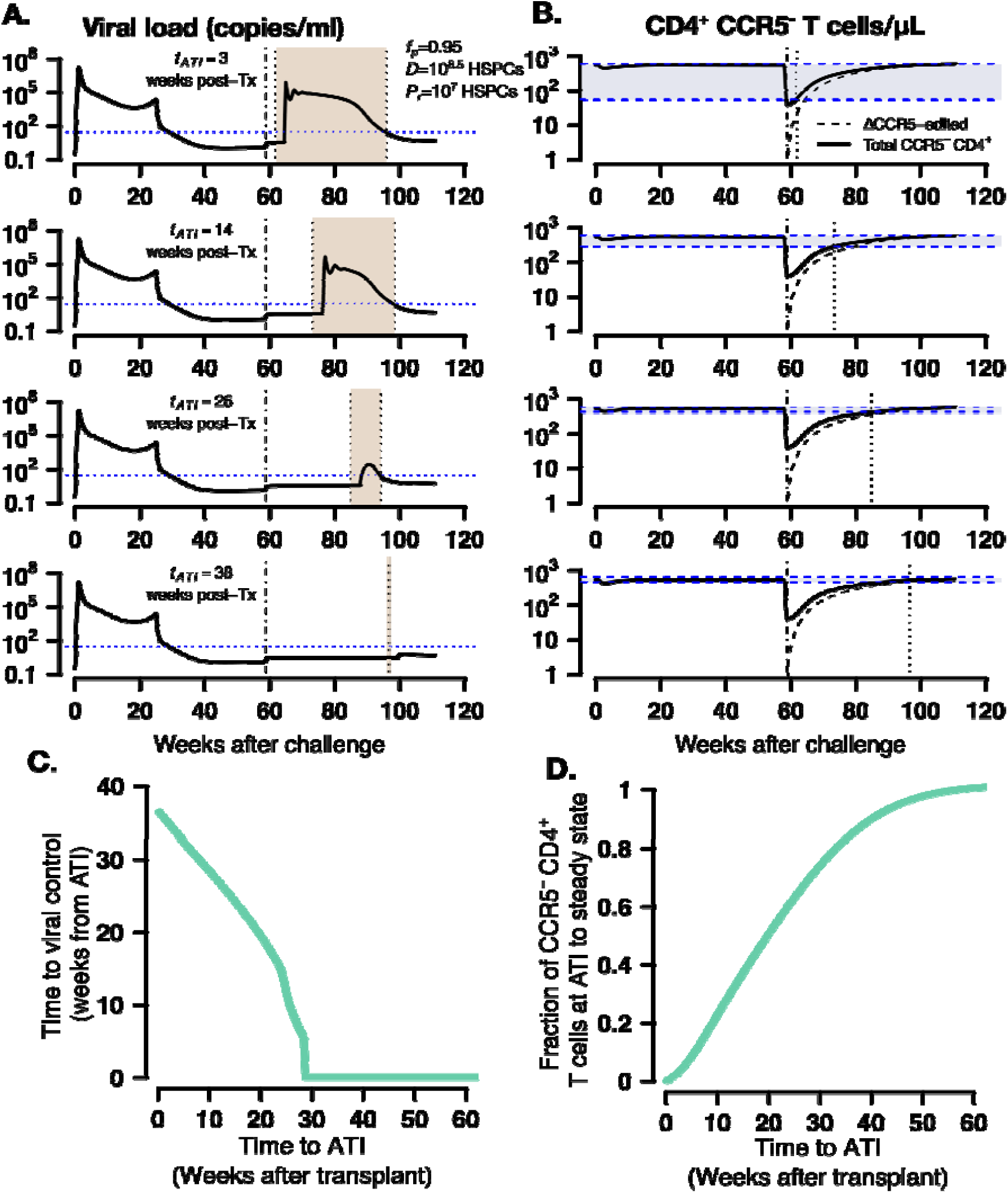
Model predictions of time to post-ATI viral control given varying times for the start of ATI. **A-B.** Examples of projected **A.** viral load and **B.** total CD4^+^ CCR5^-^ and ΔCCR5 CD4^+^ T cells from the model for animal A11219 when, HSPCs and HSPCs, for different times of ATI (= 3, 14, 26 and 38 weeks after transplantation). Dashed-dotted vertical lines represent time of transplant. Shaded areas between the dotted lines in **A.** describe the time from ATI until spontaneous viral control. Shaded areas between the blue dashed lines in **B.** represent the difference between the CD4^+^ CCR5^-^ T cell concentration at ATI and the projected steady state. Dotted lines in **B.** represent time of ATI. **C-D.** Model predictions of the **C.** time until viral control after ATI and **D.** the fraction of total CD4^+^ CCR5^-^ T cell concentration at ATI with respect to its steady state for varying times of ATI between 0 and 60 weeks after transplant for animal A11219.

In summary, our model predicts that post-ATI viral control during autologous HSPC transplantation is obtained when 1) the transplanted HSPC dose is significantly higher than the residual endogenous HSPCs that persist through myeloablative conditioning (in this case TBI) *and* 2) the fraction of protected (i.e. CCR5-edited) HSPCs in the transplant (*f*_*p*_) is sufficiently high to outcompete cells susceptible to infection and disrupt ongoing cycles of viral replication. Spontaneous post-treatment control occurs after CCR5^-^ CD4^+^ T cells achieve a steady state approximately one-year after transplantation. Hence, our model suggests that prolonging time to ATI (at least one year post-transplantation). Moreover, specifically tracking CD4^+^CCR5^-^ T cell growth and waiting for steady-state could be used as a surrogate for the decision to undergo ATI.

## Discussion

Here we introduce a data-validated mathematical model that, to our knowledge, is the first to simultaneously recapitulate viral loads as well as CD4^+^ and CD8^+^ T cell subset counts in a macaque model of suppressed HIV-1 infection. We systematically compared 12 models to arrive at a set of equations that most parsimoniously explains the available data. In multiple stages of modeling, we recapitulated (1) peripheral CD4^+^ and CD8^+^ T-cell subset reconstitution dynamics following transplant and (2) T-cell subset dynamics and SHIV viral rebound following ATI. Before ATI, all animals had suppressed plasma viral loads below the limit of detection, allowing analysis of T cell reconstitution dynamics independent of virus-mediated pressure. At each step, we applied model selection theory to select the simplest set of mechanisms capable of explaining the observed data (*37*). Our model predicts that post-rebound viral control might be possible during autologous gene-edited HSPC transplantation if therapy achieves (1) a sufficient fraction of gene-protected, autologous HSPCs, (2) a high dose of transplant product relative to a residual endogenous population of stem cells that persists following conditioning, and (3) enhancement of SHIV-specific immune responses following transplantation.

Although the model predicts a potential benefit for more potent conditioning that favors engraftment of SHIV-resistant cells, a more myeloablative conditioning regimen may also deplete SHIV-specific immune responses and lead to less favorable toxicity profiles. We previously demonstrated the link between disruption of the immune response during transplant and increased magnitude of viral rebound during treatment interruption (*13, 27*). Here we predict that the magnitude of the SHIV-specific immune response is correlated not only with viral load set point, but also with the reduction of CD4^+^CCR5^+^ T cells after ATI. CD4^+^CCR5^+^ T cell depletion might also be reflective of the loss of depletion of virus-specific immunity following conditioning. In this case, a higher threshold of CCR5-gene-edited cells in the transplanted HSPC product would be required to obtain stable, ART-free viral control. These results are consistent with the cure achieved by the Berlin patient who received a transplant with 100% HIV-resistant cells after intense conditioning (*4, 5*). In the autologous setting where 100% CCR5 editing may not be feasible, adjunctive measures that augment virus-specific immunity, such as therapeutic vaccination, infusion of HIV-specific chimeric antigen receptor (CAR) T cells or use of neutralizing antibodies, may synergize with HSPC transplantation to achieve post-treatment control (*38, 39*).

Our results are somewhat limited by a small sample size of 22 animals. For that reason, several model parameters were assumed to be the same among the population (i.e., without random effects). However, the number of observations for each animal was large enough to discriminate among several different plausible model candidates. Therefore, we performed projections using only the individual estimated parameters. Reassuringly, our results align with prior mechanistic studies of cellular reconstitution after stem cell transplantation (*15, 20, 40-42*). Our analysis also suggests that the majority of reconstituting CD4^+^CCR5^-^ T cells do not proliferate and have slow rates consistent with estimates of thymic export from previous studies (*20, 41, 42*).

Recent studies from our group and others make clear that although a preparative conditioning regimen (e.g. TBI) is essential to maximize engraftment of transplanted HSPCs, it does not clear 100% of host lymphocytes, especially those in tissues (*13, 14, 43, 44*). The best fitting model predicts that incomplete elimination of lymphocytes by TBI prevents CD4^+^CCR5^-^ cells from predominating post-transplant: the rapid expansion of CD4^+^CCR5^+^ and CD8^+^ T cells during the first few weeks after HSPC transplantation is most likely due to lymphopenia-induced proliferation of residual endogenous cells after TBI rather than thymic reconstitution. CD4^+^CCR5^-^ T cells arising from thymic export of both transplanted and remaining cells are overwhelmed by more rapidly populating CD4^+^CCR5^+^ T cells within weeks of transplantation. Going forward, we will need to identify anatomic sites and (namely viral reservoir tissues such as spleen and lymph nodes) and associated mechanisms that allow activated CD4^+^CCR5^+^ to survive conditioning.

A final important observation from our model is that CD4^+^ T cell kinetics conducive to viral control may not be reached 30 weeks after transplant. Therefore, our model suggests that ATI should be delayed until CD4^+^CCR5^-^ T cells reconstitute to their natural steady state. Optimized timing of ATI would ideally be based on reconstitution of all CD4^+^ and CD8^+^ T cell subsets ensuring approximately steady state levels before discontinuing ART.

In conclusion, our mathematical model recapitulates, to an unprecedented degree of accuracy, the complex interplay between reconstituting HIV-susceptible CD4^+^ T cells, HIV-resistant CD4^+^ T cells, infected cells, virus-specific immune cells, and replicating virus following autologous, CCR5-edited HSPC transplantation. Our results illustrate the capabilities of mathematical models to glean insight from preclinical animal models and highlight that modeling will be required to optimize strategies for HIV cure.

## Methods

### Study Design

We employed a multi-stage approach using ordinary differential equation models of cellular and viral dynamics to analyze data from SHIV-infected pig-tailed macaques that underwent autologous HSPC transplantation during ART and to find conditions for post-rebound control when gene-edited cells were included in the transplant product. First, we modeled T cell dynamics and reconstitution following transplant and before ATI, assuming that low viral loads during suppressive ART do not affect cell dynamics (**Fig. 1B**). In the second stage, we added viral load data during primary infection and after ATI and fit models to the T cell and viral dynamics simultaneously from data pre- and post-ATI (**Fig. 1C**). We then used the most parsimonious model, as determined by AIC, to perform simulated experiments for different transplant conditions, focusing on variables including fraction of protected cells, dose, depletion of HSPCs after conditioning, and time of ATI after transplant to find thresholds for viral control post-ATI.

### Experimental Data

Twenty-two juvenile pigtail macaques were intravenously challenged with 9500 TCID50 SHIV-1157ipd3N4 (SHIV-C) (*13, 14*). After 6 months, the macaques received combination antiretroviral therapy (ART): tenofovir (PMPA), emtricitabine (FTC), and raltegravir (RAL). After ∼30 weeks on ART, 17 animals received total body irradiation (TBI) followed by transplantation of autologous HSPCs. In 12/17 animals the transplant product included CCR5 gene-edited HSPCs (ΔCCR5 group); HSPC products in 5/17 animals were not edited (WT group). After an additional 25 weeks following transplant, when viral load was well suppressed, animals underwent ATI (*13*). A control group of five animals did not receive TBI or HSPC transplantation and underwent ATI after ∼50 weeks of treatment. One and six of the animals in the WT and ΔCCR5 groups, respectively, were necropsied before ATI. One of the animals in the control group was necropsied before ATI (**Fig. 1A**). Plasma viral loads and absolute peripheral T-cell counts from CD4^+^CCR5^-^, CD4^+^CCR5^+^ and total CD8^+^ and subsets (naïve, central memory [T_CM_], and effector memory [T_EM_]) were measured for the control and WT group as described previously (*13*). We analyzed peripheral T cell counts and plasma viral load from transplant until 43 weeks post-transplant (∼25 weeks pre-ATI and ∼20 weeks post-ATI).

### Mathematical modeling of T cell reconstitution dynamics

We modeled the kinetics of CD4^+^ and CD8^+^ T cell subsets in blood including residual endogenous, transplanted cells that home to the BM, and progenitor cells in the BM/thymus both from transplant and residual endogenous. We included CD8^+^ T cells in the model because CD8^+^ and CD4^+^ T cells may arise from new naïve cells from the thymus and compete with each other for resources that impact clonal expansion and cell survival (*15, 45, 46*). We assumed that expansion of CD4^+^ and CD8^+^ T cells in the blood derives from: (1) export of naïve cells differentiated from a progenitor compartment in the BM/Thymus (*40, 47*) (either from transplanted (*20, 21*) or residual endogenous CD34^+^ HSPCs) and further differentiation to an activated effector state (*19, 22-26, 48-50*), or (2) lymphopenia-induced division of mature, residual endogenous cells that persist through myeloablative TBI (*15-19*) as factors that drive T cell proliferation are more accessible (i.e., self-MHC molecules on antigen-presenting cells (*22, 23, 51*) and *γ*-chain cytokines such as IL-7 and IL-15 (*16-18, 52*)). However, as they grow, cells compete for access to these resources, limiting clonal expansion (*15*) such that logistic growth models are appropriate (*45*).

In our mathematical model, transplanted HSPCs *T* home to the bone-marrow at a rate *k*_*e*_. We assumed a single cell compartment for T cell progenitors in the bone marrow (BM)/thymus represented by variable *P*. We assumed that *P* renew logistically with maximum rate *r*_*p*_, differentiate into naïve CD4^+^ and CD8^+^ T cells at rates *λ*_*f*_ and *λ*_*e*_, respectively, or are cleared at rate *d*_*p*_ (*53-55*). We assumed two CD4^+^ T cell compartments: SHIV-non-susceptible, i.e. CD4^+^ T cells that do not express CCR5 (CD4^+^CCR5^-^ T cells) *N*, and a SHIV-susceptible compartment, *S* (CD4^+^CCR5^+^ T cells). Only the *N* compartment includes CD4^+^ naïve cells migrating from the thymus (*56-58*) at an input rate *λ*_*f*_*P* cells per day (*20, 59*). *N* cells grow with maximum rate *r*_*n*_, upregulate CCR5 (*24*) at rate *λ*_*n*_, and are cleared from the periphery at rate *d*_*n*_. The *S* compartment does not have a thymic input but can grow with maximum division rate *r*_*s*_, downregulate CCR5 (*24*) at a rate *λ*_*s*_, and are cleared at rate *d*_*s*_. We model CD8^+^ T cell reconstitution assuming a compartment for naïve and central memory cells, *M*, and a compartment for the effector memory subset, *E*. We assumed that *M* cells have thymic input of *λ*_*e*_*P* cells per day, grow logistically with maximum division rate *r*_*m*_, differentiate to effector memory at rate *λ*_*m*_, and are cleared at rate *d*_*m*_. The *E* compartment grows with maximum division rate *r*_*e*_ and is cleared at rate *d*_*e*_. We added variables *T*_*p*_, *P*_*p*_ and *N*_*p*_, representing CCR5 gene-modified-transplanted HSPCs, T cell progenitor cells in BM/thymus, and blood CD4^+^CCR5^-^ T cells, respectively. These compartments have the same structure as *T, P* and *N*, but with two differences. First, the value of *T*_*p*_ at transplantation is a fraction *f*_*p*_ of the total number of infused cells. Second, the *N*_*p*_ cell compartment do not upregulate CCR5. We model the competition of CD4^+^ and CD8^+^ T cells for resources that allow cell division using a logistic equation that depends on the difference between the total number of competing cells, i.e. *A* = *N*_*p*_+*N*+*S*+*M*+*E*, and a carrying capacity *K* (*15*). Under these assumptions we constructed the following model form:

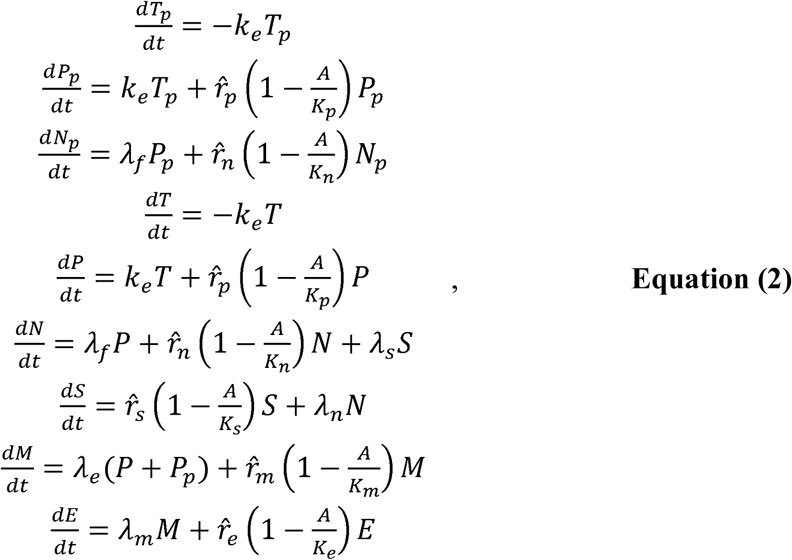

where 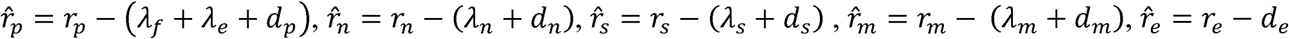, as well as 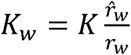 for each model variable *w* ∈ {*p, n, s, m, e*}.

When simulating the model, we assumed *t*_0_ as the time of transplantation. For the transplant groups the system is in a transient stage due to conditioning (TBI) at *t*_0_, therefore initial values cannot be obtained from steady state equations. Transplantation is modeled as *T*(*t*_0_) = (1 − *f*_*p*_) *D* and *T*_*p*_(*t*_0_) = *f*_*p*_*D.* For the control group we used *t*_0_ at a similar time relative to the transplant groups. Since the control group did not have any transplantation or TBI, we assumed *T*(*t*_0_) = *T*_*p*_(*t*_0_) = *P*_*p*_(*t*_0_) = *N*_*p*_(*t*_0_) = 0. Other initial values were calculated assuming steady state: 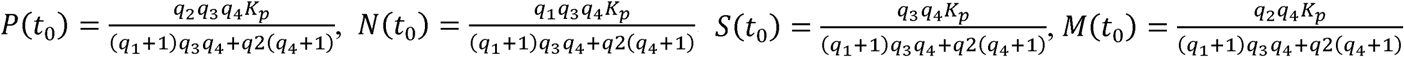 and 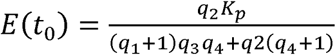. Here 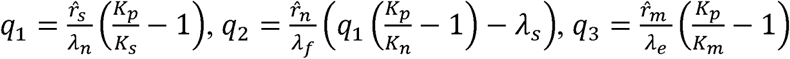 and 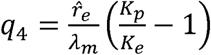. A parsimonious, curated version of this model was selected from a series of models with varying mechanistic and statistical complexity (**Table S2).**

### Mathematical modeling of SHIV infection and T cell response dynamics

We next adapted the curated T cell reconstitution model by combining several adaptations of the canonical model of viral dynamics (*27, 60-66*). Here, virus *V* infects only CD4^+^CCR5^+^ T cells (*67*) *S* at rate *β*. We modeled ART by reducing the infection rate to zero. A fraction *τ* of the infected cells produce virus, *I*_*p*_, and the other fraction become unproductively infected, *I*_*u*_ (*27, 68, 69*). *I*_*P*_ cells arise only from activation of a persistent set of latently infected cells at rate 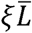. We modeled ATI by assuming infection *β* is greater than zero after some delay following ART interruption. We approximate this delay as the sum of the time of ART to washout (∼3 days) and the time of successful activation (*t*_*sa*_) of a steady set of latently infected cells. For simplicity, we assumed that 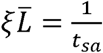 and *t*_*sa*_ assumed that has lognormal distribution among the animal population with high variance (*70-72*). All infected cells die at rate *δ*_*I*_ (*27*). *I*_*P*_ cells produce virus at a rate *π* per cell, that is cleared at rate *γ*. CD8^+^ *M* cells proliferate in the presence of infection with maximum rate *ω*_*8*_. A fraction *f* of these cells become SHIV-specific CD8^+^ effector T cells, *E*_*h*_, that are removed at a rate *d*_*h*_ (*64, 65, 73*). These effector cells may reduce virus production (*π*) or increase infected cell clearance (*δ*_*I*_) by 1/(1+θ*E*_*h*_) or by (1+κ*E*_*h*_), respectively (*28-30, 63, 74*). We assumed that non-susceptible CD4^+^ T cells may upregulate CCR5 and replenish the susceptible pool during infection (*35, 36, 75*) with rate *ω*_*4*_. For cell growth the total number of competing cells is given by *A* = *N*_*p*_+*N*+*S*+*I*_*p*_+*I*_*u*_+*M*+*E*+*E*_*h*_. The model in **eq. 2** is modified to include:

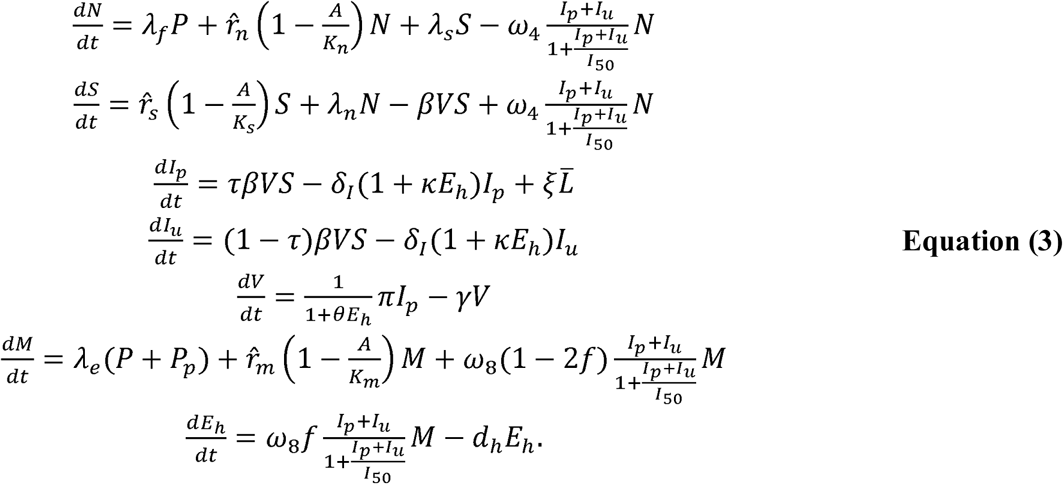

When simulating this model, we assume *t*_0_ = 0 as the moment of SHIV challenge, and *t*_*x*_ as the moment of transplantation after challenge. We modeled conditioning by: (1) adding a term −*k*_*T*_*C* in all blood cell compartments *C* ∈{*N*_*p*_,*N,S,I*_*p*_,*I*_*u*_,*M,E,E*_*n*_} and (2) the term −*k*_*H*_*P* for the HSPC compartment *P. k*_*T*_ and *k*_*H*_ are different than zero only during the two days before transplant (*t*_*x*_ − 2≤ *t* < *t*_*x*_). Transplantation is modeled as an input only when *t* = *t*_*x*_ to cell compartments *T* and *T*_*p*_ with amounts (1 − *f*_*p*_)*D* and *f*_*p*_ *D*, respectively. A parsimonious version of this model was selected from a series of models with varying mechanistic and statistical complexity (**Table S2).**

### Fitting procedure and model selection

To fit our models (**eqs. 2, 3**) to the transplant data, we used a nonlinear mixed-effects modeling approach (*76*) described in detail in the **Supplementary Materials**. Briefly, we obtained a maximum likelihood estimation of the population median (fixed effects) and standard deviation (random effects) for each model parameter using the Stochastic Approximation Expectation Maximization (SAEM) algorithm embedded in the Monolix software (www.lixoft.eu). For a subset of parameters, random effects were specified, and the standard deviation values were estimated. Measurement error variance was also estimated assuming an additive error model for the logged outcome variables.

We first fit instances of models with varying statistical and mechanistic complexity in **eq. 2** to blood T cell counts during transplant and before ATI (**Fig. 1B**) assuming that one or multiple mechanisms are absent, or that certain mechanisms have equal kinetics (**Table S2** includes all 12 competing models with the different statistical assumptions). We fixed the HSPC homing rate *k*_*e*_ = 1/day (*77, 78*), and *f*_*p*_ and *D* as described in **Table S1**. Since at (*t*_0_) the system is in a transient stage due to conditioning (TBI) we estimated blood cell concentrations at (*t*_0_), but fixed the number of HSPCs that remained in the bone marrow/thymus *P*(*t*_0_) to 6×10^6^ based on the estimated minimum number of infused HSPCs needed for engraftment in the same animal model (*44*). All other parameters were estimated were estimated as described in **Supp. Materials** and **Table S2**.

Next, we fit several instances of the model in **eq. 3** to pre- and post-ATI blood T cell counts and plasma viral loads (**Fig. 1B**) using the best model obtained for **eq. 2** (**Table S5** includes all 4 competing models and respective statistical assumptions). At the time of SHIV infection, values for the cell compartments were calculated from steady state equations with the same form as for the group without transplantation (“control”) in the previous section. *V*(0) was fixed to a small value below the limit of detection, and *I*_*p*_(0) and *I*_*u*_(0) were calculated as *τcV*(0)/*π* and (1 − *τ*) *cV*(0)/*π*, respectively. We fixed the following parameters: *γ* = 23/day (*79*), *δ*_*I*_ = 1/day (*80, 81*), *τ* = 0.05 (*68*) and *f* = 0.9 (*63*). The value of *k*_*h*_ was constrained to obtain a value of the HSPCs after conditioning *P*(*t*_*x*_) = *P*_*r*_ = 6 × 10^6^ (*44*). We fixed values of *t*_*x*_, *f*_*p*_ and *D* as described in **Table S1.** All other parameters were estimated as described in **Supp.Materials** and **Table S5**.

To determine the best and most parsimonious model among the instances, we computed the log-likelihood (log *L*) and the Akaike Information Criteria (AIC=-2log *L*+2*m*, where *m* is the number of parameters estimated) (*37*). We assumed a model has similar support from the data if the difference between its AIC and the best model (lowest) AIC is less than two (*37*).

### *Effective reproductive ratio when r*_*n*_ = 0 *and k* = 0

We calculated an approximate effective reproductive ratio *R*_*eff*_ for our model (**eqs. 2, 3**) by computing the average number of offspring produced by one productively infected cell *I*_*p*_ at ATI assuming all cell compartments have reached steady state after transplantation during ART. This number is the product of the average lifespan of one *I*_*p*_, the virus production rate by this latently infected cell, the lifespan of produced virions from this cell, the rate at which each virion infects the pool of susceptible cells at steady state, the fraction of these infections that become productive and the reduction of virus production, cell infection and cell death by SHIV-specific immune cells at ATI. Using this approach we obtain that 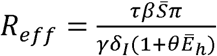, with 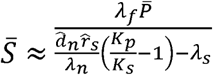 and 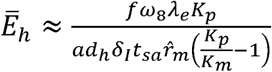 the steady state values of variables *S* (SHIV-susceptible cells) and *E*_*h*_ (SHIV-specific effector cells) during ART with 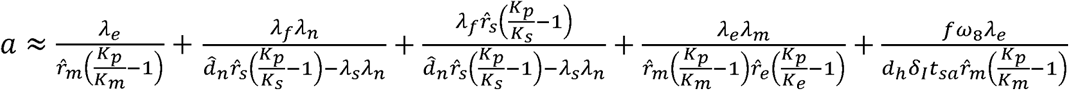. By assuming that the total amount of infused cells (dose *D* and fraction of CCR5-editing *f*_*p*_) home to the BM/Thymus rapidly, and that the amount of remaining HSPCs after TBI and immediately before transplant is *P(t*_*x*_) = *P*_*r*_, the approximate steady state for *P* is 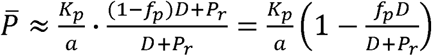. Together this gives the following expression for the effective reproductive ratio:

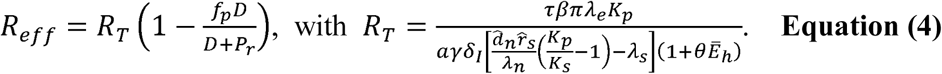

Here, *R*_*T*_ then represents the effective reproductive ratio during transplant in the absence of gene-editing when cells have reached steady state.

### *Approximate viral load set point when r*_*n*_ = 0 *and k* = 0

We calculated an approximate viral steady state after ATI assuming all derivatives in **eq. (3)** are equal to zero. This assumption leads to the equations, 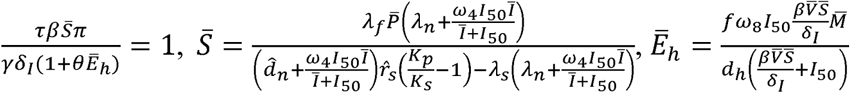 and 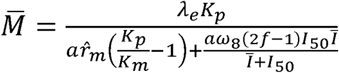. To obtain an approximation of the viral set point 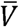 we used the equations above and the following rules:

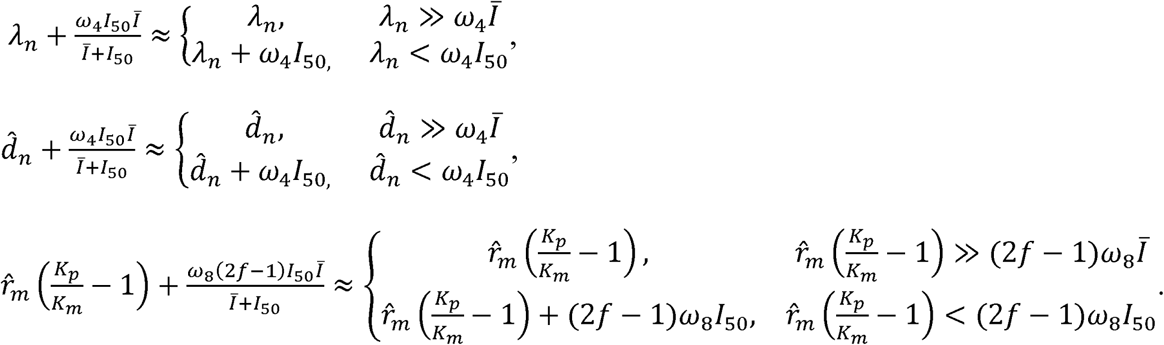

Here, *Ī* is obtained from simulations of the model after 10 years after ATI for *I*_*p*_ + *I*_*u*_ using parameter estimates from best fits to each animal data. Regardless of the assumptions above, the viral load set point is always proportional to the following term,

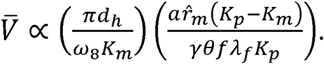

We define the ratio 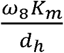 as the turnover rate of SHIV-specific CD8^+^ T cells.

## Supplementary Materials

### Supplementary Methods

#### Supplementary Figures

Fig. S1. CD4^+^ and CD8^+^ T cell levels pre-ATI in control group (n = 5) at times relative to post-transplantation in WT and ΔCCR5 transplant groups.

Fig. S2. Individual fits of the best model to the blood T cell observations pre-ATI in control group from a time relative to post-transplantation in transplant groups.

Fig. S3. Individual fits of the best model to the blood T cell observations post-transplantation, pre-ATI for the wild-type-transplant group.

Fig. S4. Individual fits of the best model to the blood T cell observations post-transplantation, pre-ATI for the ΔCCR5-transplant group.

Fig. S5. Predictions of the best model for the contributors to cell expansion in CD8+ T_EM_ cells in animals from the transplant groups.

Fig. S6. Correlations between viral load and CD4^+^CCR5^+^ T cells over time in each animal post-ATI.

Fig. S7. Individual fits of the best model to the blood T cell and viral load observations before/after ATI for control group.

Fig. S8. Individual fits of the best model to the blood T cell and viral load observations before/after ATI for the wild-type-transplant group.

Fig. S9. Individual fits of the best model to the blood T cell and viral load observations before/after ATI for the ΔCCR5-transplant group.

Fig. S10. Model predictions for post-rebound viral control after CCR5 gene-edited HSPC transplantation based on *R*_*eff*_.

Fig. S11. Model predictions of the fraction of protected HSPCs in the transplant *f*_*p*_ (y-axis) and the fraction of transplanted HSPCs with respect to the total infused plus remaining post-TBI HSPCs *D*: *P*_*r*_ (x-axis in log-scale) required for spontaneous viral axis) control.

#### Supplementary Tables

Table S1. Values of the fraction of protected cells in transplant product *f*_*p*_, dose or number of HSPCs in transplant product *D* and time of transplantation *t*_*x*_ of each animal for model fitting and projections.

Table S2. Competing models for fitting T cell reconstitution with respective AIC values.

Table S3. Population parameter estimates for the best fits of the model in eq. 2 in the main text (lowest AIC in Table S2) to the T cell reconstitution dynamics.

Table S4. Individual parameter estimates for the best fits of the model in eq. 2 in the main text (lowest AIC in Table S2) to the T cell reconstitution dynamics.

Table S5. Competing models for fitting T cell and viral dynamics (eqs. 2-3 in main text) using the best model in Table S2 and fixing parameter values as in Table S3, with AIC values.

Table S6. Population parameter estimates for the fits of the model with lowest AIC in Table S5 to the T cell and virus dynamics.

Table S7. Individual parameter estimates for the fits of the model in eqs. 2-3 in main text (lowest AIC in Table S5) to the T cell and virus dynamics.

## Author contributions

E.F.C. and J.T.S. conceived the study. C.W.P. and H.-P.K. contributed ideas and data sources for the project. E.R.D. and D.B.R. contributed in the development of mechanistic mathematical models. B.T.M. contributed ideas and support for statistical models and analyses. E.F.C. assembled data, wrote all code, performed all calculations and derivations, ran the models, and analyzed output data. J.T.S. and E.F.C. wrote the manuscript with contributions from all other authors.

## Animal Welfare

The data used in this work were collected in strict accordance with the recommendations in the *Guide for the Care and Use of Laboratory Animals* of the National Institutes of Health. The study protocol was approved by the Institutional Animal Care and Use Committees (3235-03) of the Fred Hutchinson Cancer Research Center and the University of Washington.

## Data and materials availability

Original data and code will be shared upon request.

