## Supplementary Materials for "Thresholds for post-rebound SHIV control after CCR5 gene-edited autologous hematopoietic cell transplantation"

**Authors:** E. Fabian Cardozo-Ojeda^1^, Elizabeth R. Duke^1,4^, Christopher W. Peterson^2,3,4^, Daniel B. Reeves^1^, Bryan T. Mayer^1^, Hans-Peter Kiem^2,3,4,5^, Joshua T. Schiffer^1,2,4,*^

**Affiliations:**

^1^Vaccine and Infectious Disease Division,

^2^Clinical Research Division,

^3^Stem Cell and Gene Therapy Program, Fred Hutchinson Cancer Research Center, Seattle, WA, USA;

^4^Department of Medicine and

^5^Department of Pathology, University of Washington, Seattle, WA, USA.

### Supplementary Methods

### 1. Notes on nonlinear, mixed-effects modeling

To model the longitudinal T cell and viral load observations for all animals we used a nonlinear, mixed-effects modeling approach. Within this approach we modeled a state variable vector v with observations at time$i$ for each animal $j$ as,

$v_{ij}=f_{v}\left( t_{ij}{,\Psi}_{j} \right)+\epsilon_{v}$,	**(S1)**

where $f_{v}$ is a *nonlinear* function for the state variable vector $v$ at an individual observation time, $t_{ij}$, with animal-specific parameter set $\Psi_{j}$. The distribution of measurement noise, $\epsilon_{v}$, is normally distributed with a state-variable-specific standard deviation $\sigma_{v}$,

$\epsilon_{v}\mathcal{\sim N}\left( 0,\sigma_{v}^{2} \right)$,	**(S2)**

In the *mixed-effects* model it is assumed that for an animal $j$ each single parameter ${\psi_{j}\in\Psi}_{j}$is drawn from a probability distribution across the population. This distribution includes the fixed effects $\bar{\psi}$ representing the median value over the population, and the random effects $\eta_{j}$ representing its variability in the population, assumed to be normally distributed with standard deviation $\sigma_{\psi}$, that is

$\eta_{j}\mathcal{\sim N}\left( 0,\sigma_{\psi}^{2} \right)$.	**(S3)**

We assumed that the random effects of the parameters$\eta_{j}$ might not be independent. In that case the vector of random effects $\eta_{j}$follows a multinormal distribution: $\eta\mathcal{\sim N}\left( 0,\Omega\right)$, being $\Omega$ the variance-covariance matrix based on the values $\sigma_{\psi}$ and correlations between the individual parameters in $\eta$_._

The non-linear function, $f_{v}$, for the given state variables are estimated using numerical solutions of the differential equation models described in the main text (**eqs. 2,3**). We fit each model to all data points from all animals simultaneously using a maximum likelihood approach. We assumed that individual observations of each state variable $v_{ij}$ for each animal $j$ at each time point $t_{ij}$ are independent. For each model we obtained the Maximum Likelihood Estimation (MLE) of the standard deviation of the measurement error for the observations $\sigma_{v}$, and each parameter fixed effects $\bar{\psi}$ and standard deviation of the random effects $\sigma_{\psi}$ (or elements in matrix $\Omega$ when applicable) using the Stochastic Approximation of the Expectation Maximization (SAEM) algorithm embedded in the Monolix software (www.lixoft.eu)^1^.

### 2. Notes on fitting T cell reconstitution before ATI

We first fit the observed blood T cell kinetics after hematopoietic stem and progenitor cell (HSPC) transplantation and before analytical treatment interruption (ATI). During this procedure, we defined the vector for the state variables $v^{(1)}$ as

$v^{(1)}=\{S,N,C,M,E\}$	**(S4)**

representing the observed blood CD4^+^CCR5^-^, CD4^+^CCR5^+^, total CD8^+^, CD8^+^ T_N_ + T_CM_, and CD8^+^ T_EM_ cell counts, respectively. We modeled the kinetics of $v^{(1)}$ using the nonlinear ordinary differential equation (ODE) system in **eq. 2** in the main text, with solution $f^{(1)}$. In this model we assumed that the total number of CD8+ T cells is defined as $C=M+E$.

We defined the statistical form of each parameter in $\Psi^{(1)}$ by using different functional forms. In complete detail: parameters $\hat{r}_{p}^{j}, \hat{r}_{m}^{j},\hat{r}_{e}^{j},\lambda_{f}^{j},\lambda_{e}^{j},\lambda_{n}^{j},\lambda_{s}^{j},\lambda_{m}^{j}$ were modeled as $\psi_{j}=\bar{\psi}e^{n_{j}}$; parameter $K_{p}^{j}$ was modeled as $\psi_{j}={10}^{{\bar{\psi}+n}_{j}}$; $K_{n}^{j},K_{s}^{j},K_{m}^{j},K_{e}^{j}$ were modeled as $\psi_{j}={10}^{K_{p}^{j}-\bar{\psi}e^{n_{j}}}$; and initial values in the transplant group: $N^{j}\left( t_{0} \right),S^{j}\left( t_{0} \right),M^{j}\left( t_{0} \right)$and $E^{j}\left( t_{0} \right)$ had the model $\psi_{j}={10}^{{\bar{\psi}+n}_{j}}.$ We explored the possibility that $r_{n}=0$, in that case we assumed $\hat{d}_{n}^{j}=\lambda_{n}^{j}\left( 1+\bar{\psi}e^{n_{j}} \right)$.

We listed different competing instances of the model in **eq. 2** in the main text with different mechanistic and statistical assumptions as presented in **Table S2.**

### 3. Notes on fitting T cell and Viral load dynamics before and after ATI

For the fits of the T cell and viral dynamics after ATI we added a model for the observed plasma viral load $V$ and we also assumed that CD4^+^CCR5^+^ T cells include infected cells. Now, we define the vector for the state variables $v^{(2)}$ as

$v^{(2)}=\left\{ R,N,C_{4},C_{8},M,E,V \right\}$,	**(S5)**

with $V$ indicating the observed plasma viral load, $R$ indicating the observed blood CD4^+^CCR5^+^ T cell concentration, $C_{8}$ the total CD8^+^ T cell concentration, $C_{4}$ the total CD4^+^ T cell concentration and the others state-variables as specified for $v^{(1)}$. We included $C_{4}$ because we had total CD4^+^ T cell counts, but CD4^+^ T subset counts during the primary infection stage were not available in many of the animals. We modeled the kinetics of $v^{(2)}$ by using the ODE system in **eqs 2-3** in the main text. From this model we defined the total number of CD4^+^CCR5^+^ T cells as $R=S+I_{p}+I_{u}$ and the total number of CD4^+^ T cells as $C_{4}=R+N$.

For this model we defined the parameter set $\Psi^{(2)}$ by adding to the parameters in the previous section the parameters relative to virus dynamics (i.e.,$\Psi^{(2)}=\left\{ \Psi^{\left( 1 \right)},\kappa^{j},\theta^{j}, \beta^{j},\pi^{j}, \omega_{4}^{j}, \omega_{8}^{j}, I_{50}^{j},d_{h}^{j},t_{sa}^{j} \right\}$ but fixing the values in $\Psi^{\left( 1 \right)}$ to the MLE values using **Table S3**. For parameters $\kappa^{j},\theta^{j}, \beta^{j},\pi^{j}, \omega_{4}^{j}, \omega_{8}^{j}, I_{50}^{j}$ we used a model with form $\psi_{j}={10}^{{\bar{\psi}+n}_{j}}$, and for $d_{h}^{j}$ and $t_{sa}^{j}$ we used $\psi_{j}=\bar{\psi}e^{n_{j}}$. We evaluated single or combination of mechanistic hypotheses along with different statistical assumptions as listed in **Table S5** using AIC.

### Supplementary Figures

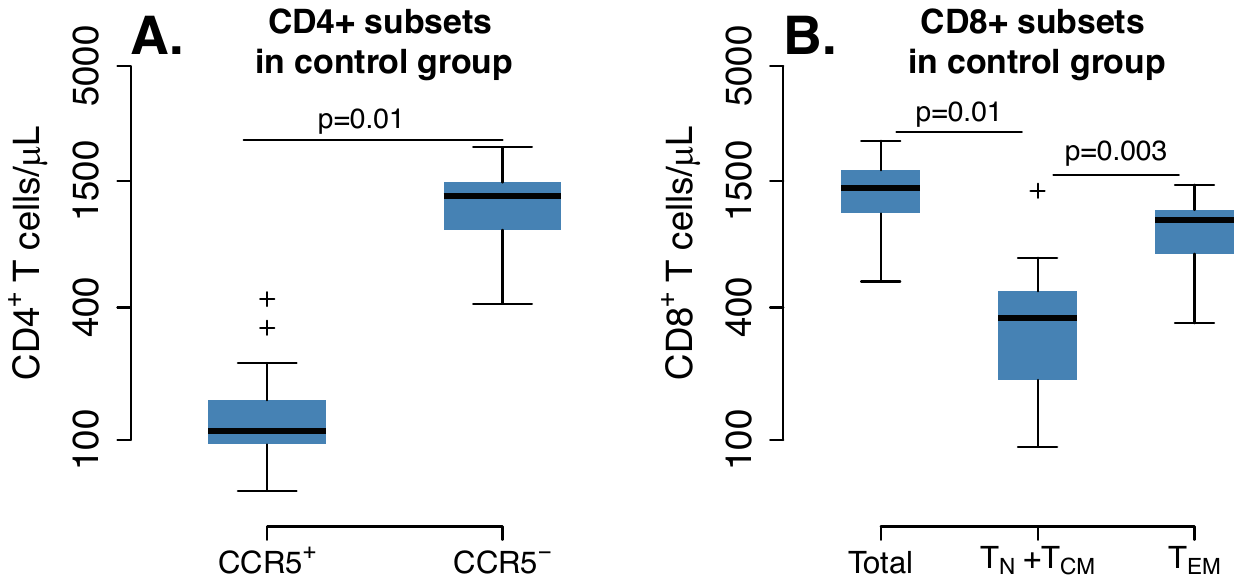

**Figure S1.** **CD4^+^ and CD8^+^ T cell levels pre-ATI in control group (n = 5) at times relative to post-transplantation in WT and ΔCCR5 transplant groups.** Range of blood **A.** CD4^+^ and **B.** CD8^+^ T cell counts using all data points for the period before ATI in control animals (p-value calculated with a paired t-test for averaged measurements from a time relative to infusion in transplanted animals and before ATI).

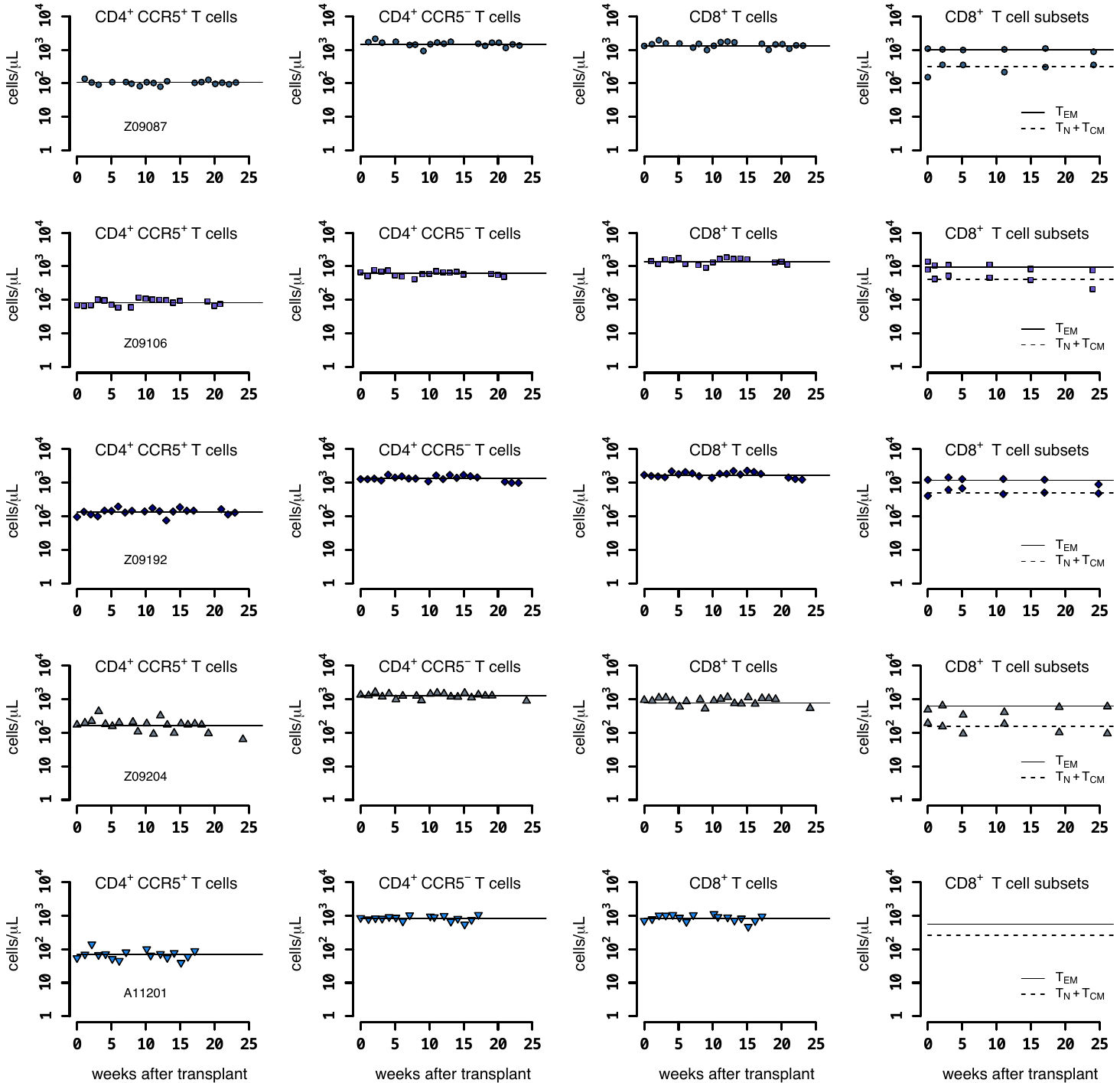

**Figure S2. Individual fits of the best model to the blood T cell observations pre-ATI in control group from a time relative to post-transplantation in transplant groups.** Empirical data for peripheral T cell subset counts (blue data points) and best fits of the model (black lines) in **eq. 2** in the main text to all blood T cell subsets before/after ATI for the control group. Each row is one animal (ID in the leftmost graph per row). Each datapoint shape and color is a different animal sampled over time and is maintained throughout.

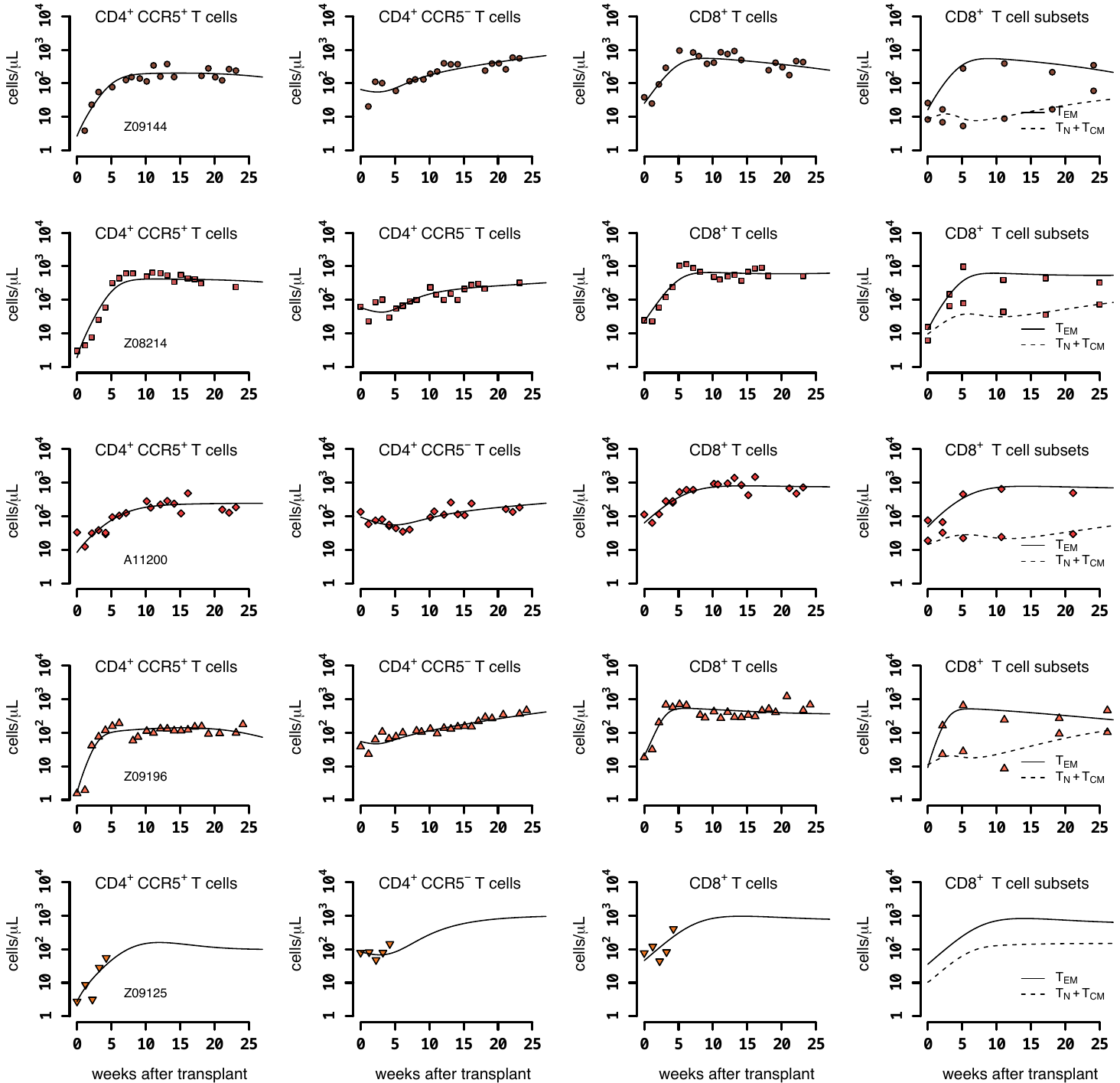

**Figure S3. Individual fits of the best model to the blood T cell observations post-transplantation, pre-ATI for the wild-type-transplant group.** Empirical data for peripheral T cell subset counts and plasma viral load (red data points) and best fits of the model (black lines) in **eq. 2** in the main text to all blood T cell subsets before ATI for the wild-type-transplant group. Each row is one animal (ID in the leftmost graph per row).

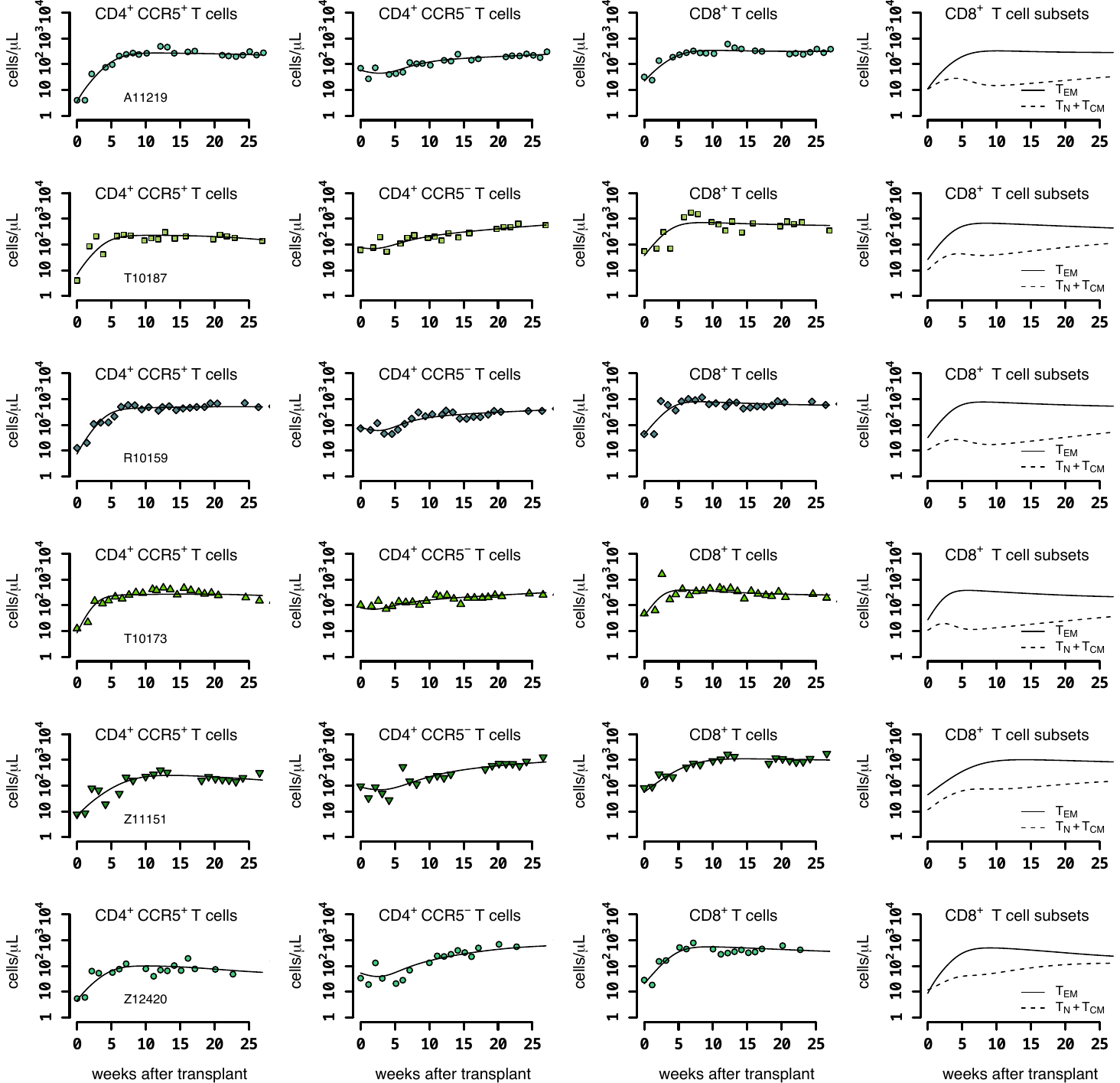

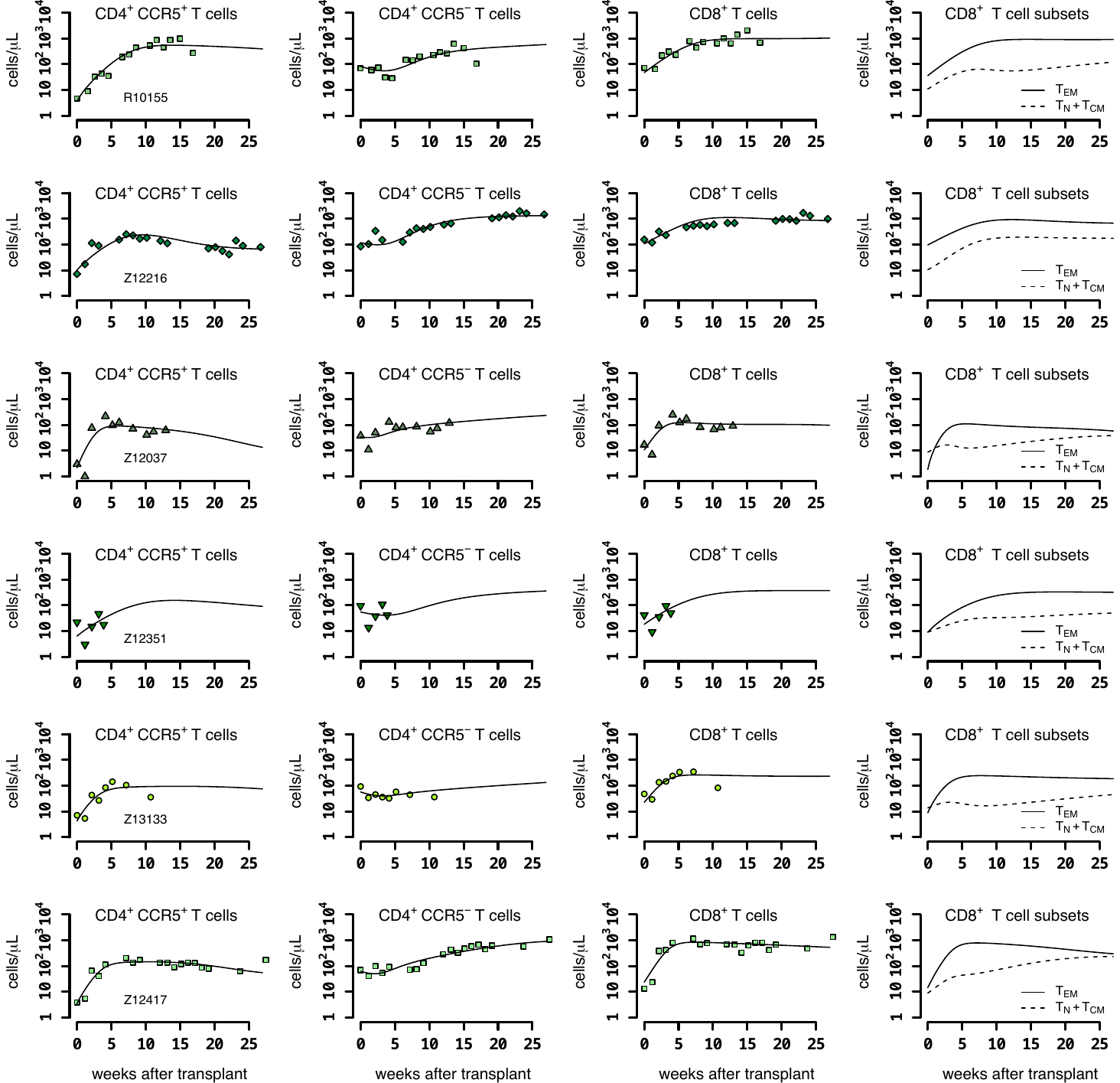

**Figure S4. Individual fits of the best model to the blood T cell observations post-transplantation, pre-ATI for the ΔCCR5-transplant group.** Empirical data for peripheral T cell subset counts and plasma viral load (green data points) and best fits of the model (black lines) in **eq. 2** in the main text to all blood T cell subsets before ATI for the ΔCCR5-transplant group. Each row is one animal (ID in the leftmost graph per row).

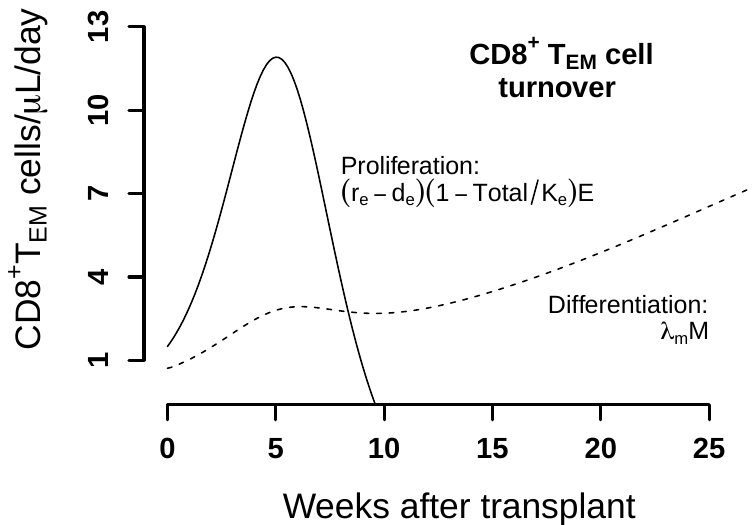

**Figure S5. Predictions of the best model for the contributors to cell expansion in CD8+ TEM cells in animals from the transplant groups.** Solid line represents the total number of cells that proliferate over time $\hat{r}_{e}\left( 1-\frac{N_{p}+N+S+M+E}{K_{e}} \right)E$. Dashed lines indicate the number of exogenous cells differentiated from T_naive_ and T_CM_ ($\lambda_{m}M$) over time using the maximum likelihood estimation of the population parameters.

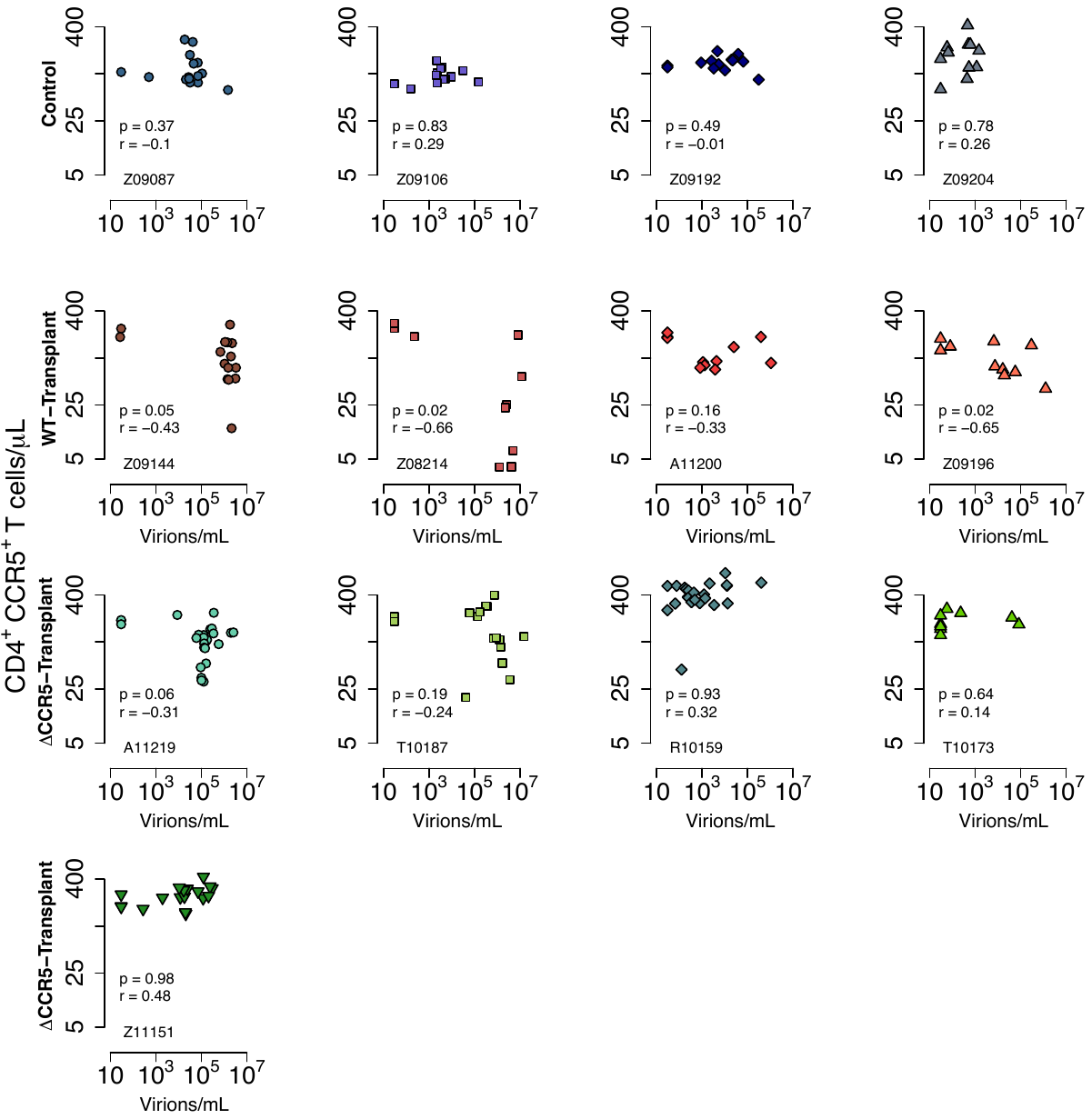

**Figure S6.** **Correlations between viral load and CD4^+^CCR5^+^ T cells over time in each animal post-ATI.** Each panel shows the timepoints post-ATI for each animal. P-values were calculated using Pearson’s correlation test Animal Z12420 in the ΔCCR5-group did not have enough detectable viral load measurements after ATI to compute the correlation and is not presented here. Animals that were necropsied before ATI are not presented here.

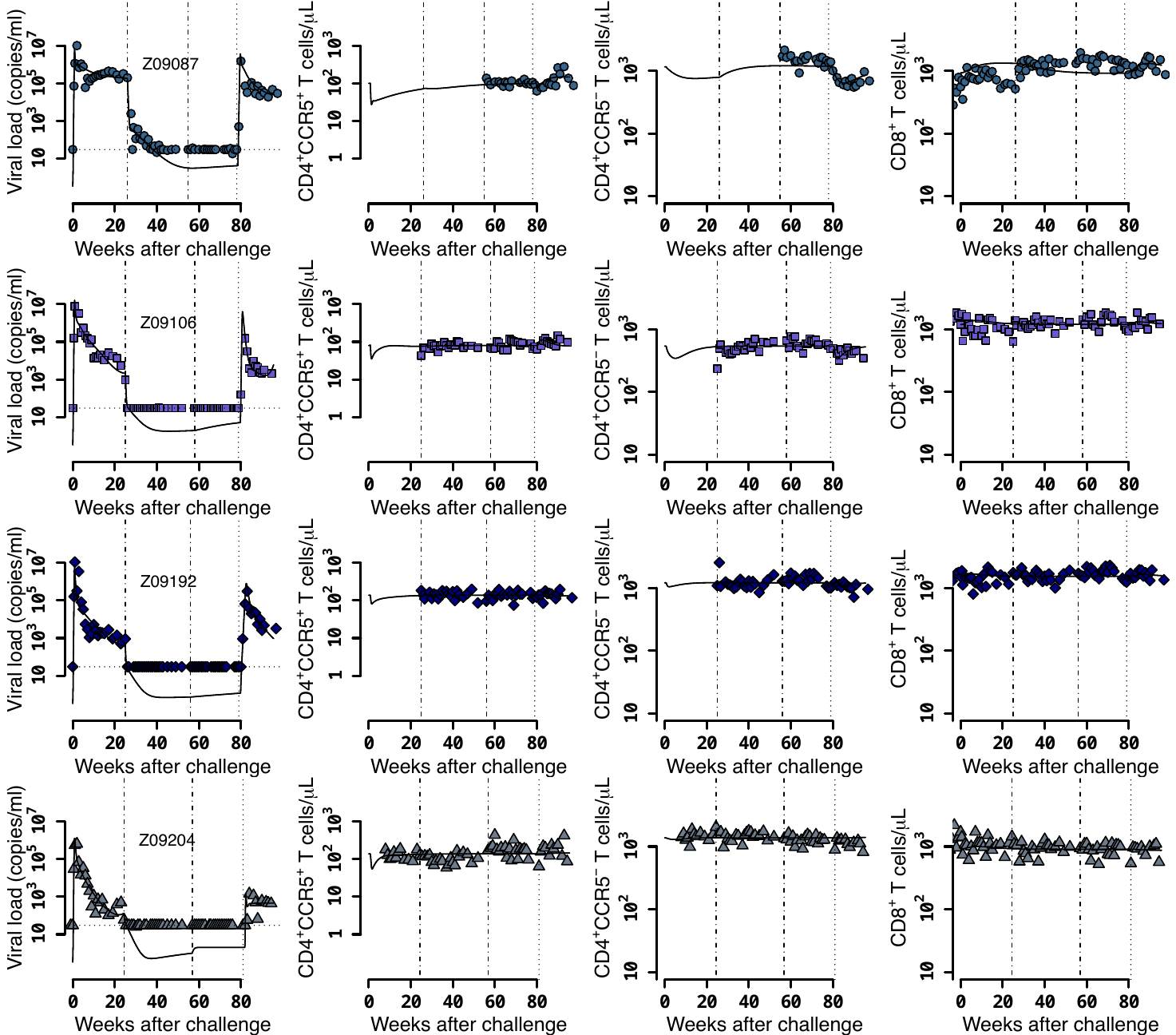

**Figure S7.** **Individual fits of the best model to the blood T cell and viral load observations before/after ATI for control group.** Empirical data for peripheral T cell subset counts and plasma viral load (blue data points) and best fits of the model in **eqs. 2 and 3** to all blood T cell subsets before/after ATI for the control group. Dashed-dot lines: cART initiation and time relative to transplantation with respect to the other groups; dotted line: ATI. Each row is one animal (ID in the leftmost graph per row).

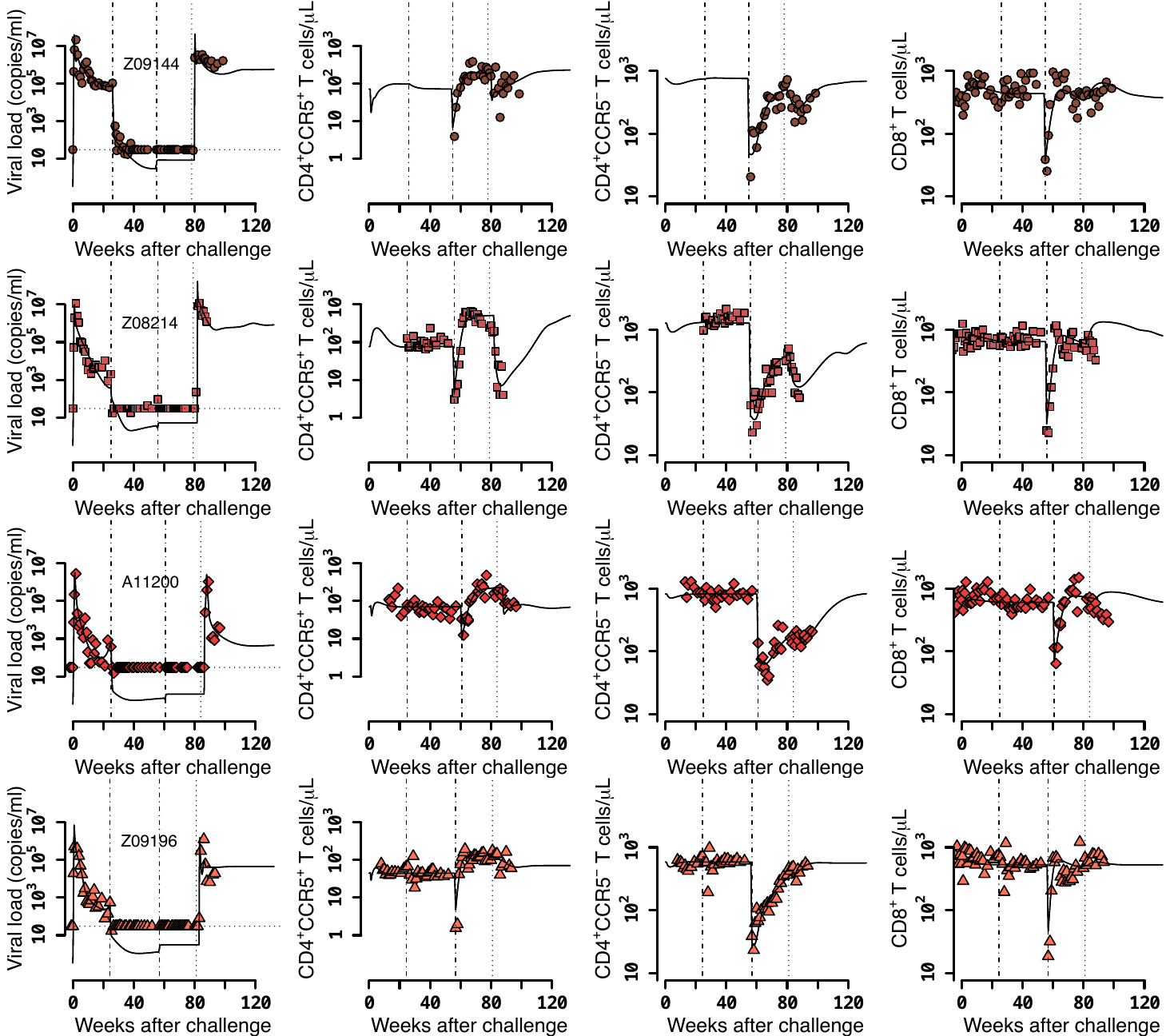

**Figure S8.** **Individual fits of the best model to the blood T cell and viral load observations before/after ATI for the wild-type-transplant group.** Empirical data for peripheral T cell subset counts and plasma viral load (red data points) and best fits of the model in **eqs. 2 and 3** to all viral load observations and blood T cell subsets before/after ATI for the wild-type-transplant group (solid lines). Dashed-dot lines: cART initiation and transplantation; dotted line: ATI. Each row is one animal (ID in the leftmost graph per row).

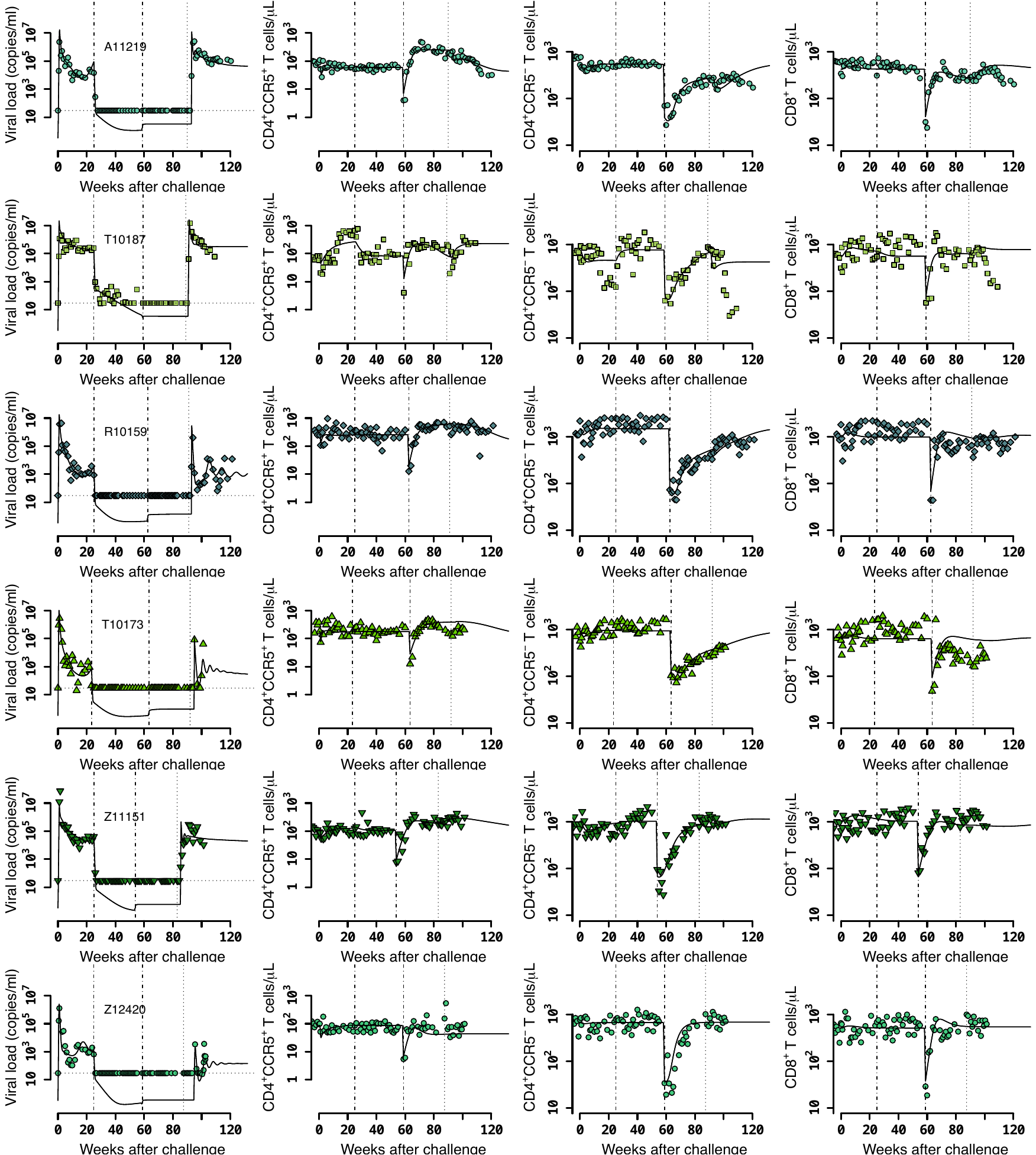

**Figure S9.** **Individual fits of the best model to the blood T cell and viral load observations before/after ATI for the ΔCCR5-transplant group.** Empirical data for peripheral T cell subset counts and plasma viral load (green data points) and best fits of the model in **eqs. 2 and 3** to all viral load observations and blood T cell subsets before/after ATI for the ΔCCR5-transplant group (solid lines). Dashed-dot lines: cART initiation and transplantation; dotted line: ATI. Each row is one animal (ID in the leftmost graph per row).

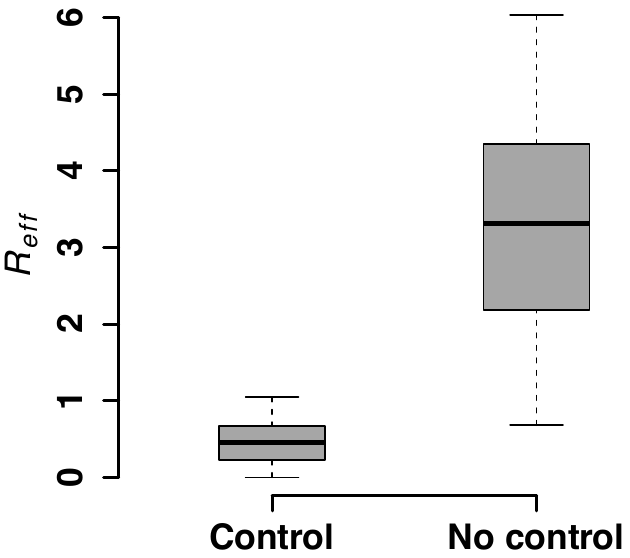

**Figure S10.** **Model predictions for post-rebound viral control after CCR5 gene-edited HSPC transplantation based on** $\boldsymbol{R}_{\boldsymbol{eff}}$**.** Model predictions of the effective reproductive ratio $R_{eff}=R_{T}\left( 1-\frac{f_{p}D}{D+P_{r}} \right)$that lead to post-ATI viral control or not. $R_{eff}$ was computed using varying values of $f_{p}$: fraction of HSPCs in transplant, $D$: total amount of infused HSPCs and $P_{r}$: remaining number of HSPCs after TBI before transplant and using parameter estimates from all animals (**Table S7**) to estimate $R_{T}$ from **eq. 4** in the main text.

| 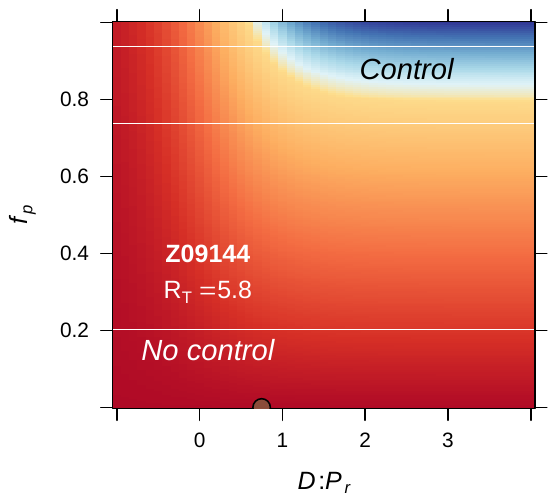 | 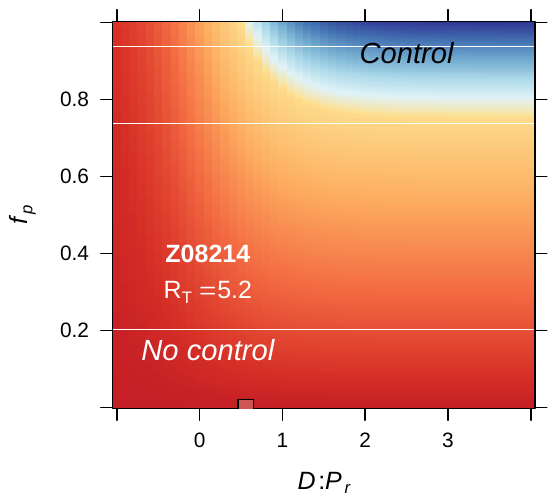 | 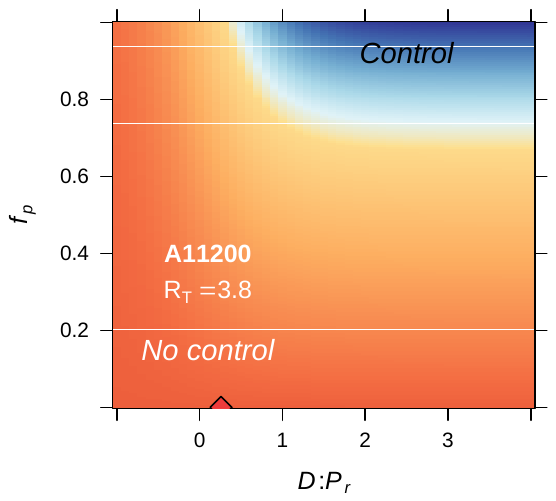 | 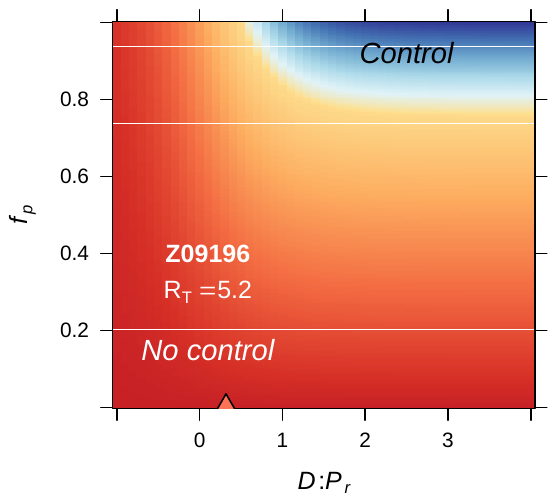 |
| --- | --- | --- | --- |
| 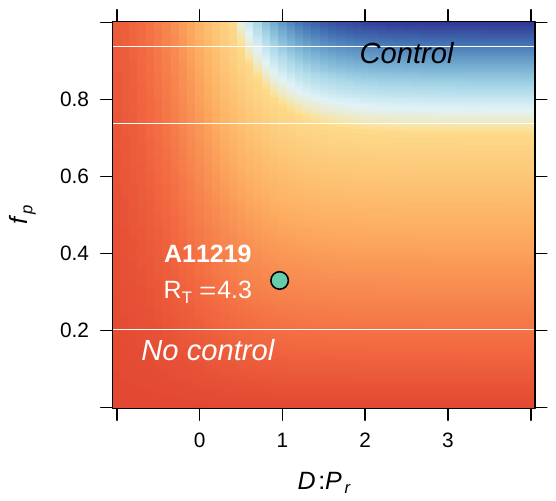 | 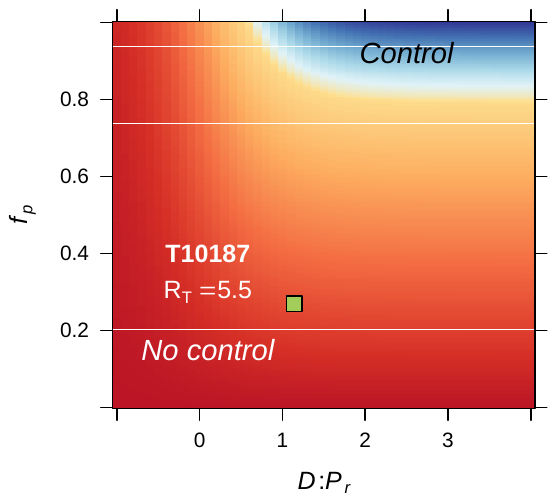 | 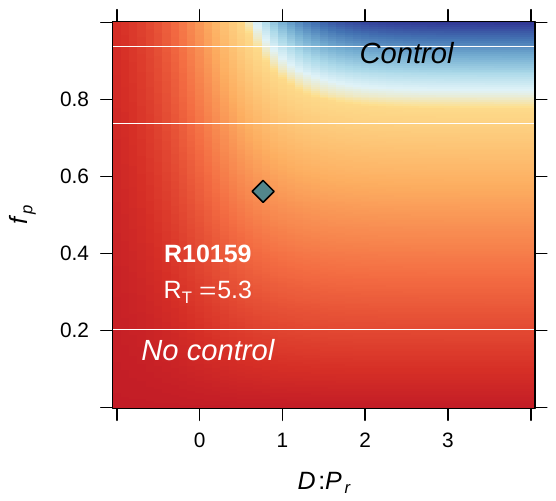 | 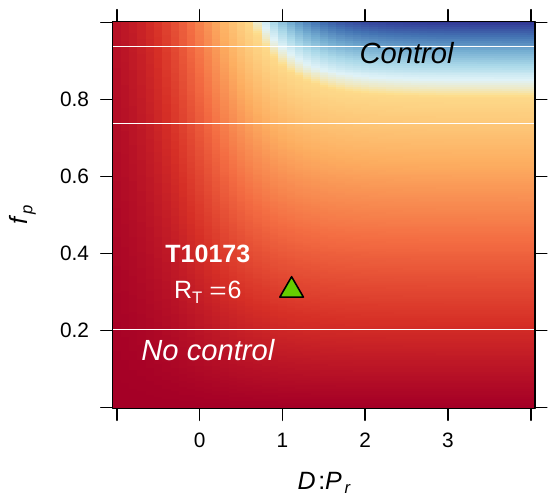 |
| 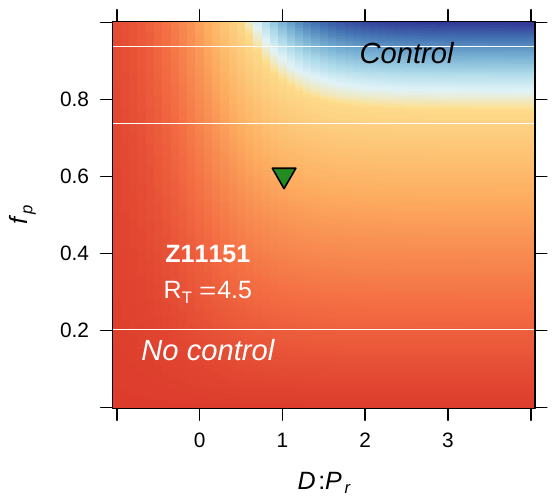 | 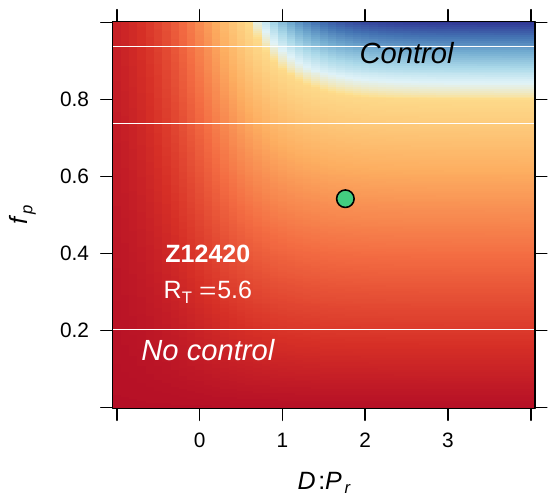 |  |  |

**Figure S11.** **Model predictions of the fraction of protected HSPCs in the transplant** $\boldsymbol{f}_{\boldsymbol{p}}$ **(y-axis) and the fraction of transplanted HSPCs with respect to the total infused plus remaining post-TBI HSPCs** $\boldsymbol{D:}\boldsymbol{P}_{\boldsymbol{r}}$ **(x-axis in log-scale) required for spontaneous viral control.** Blue color represents the parameter space with post-ATI viral control or $R_{eff}<1$. Yellow-to-red colors represent the parameter space with no control or $R_{eff}>1$. Data points (green and red shapes) represent the individual values of $f_{p}$ and $D:P_{r}$ from each transplanted animal in the study, assuming $P_{r}={10}^{7}$ HSPCs.

### Supplementary Tables

**Table S1.** Values of the fraction of protected cells in transplant product $f_{p}$, dose or number of HSPCs in transplant product $D$ and time of transplantation $t_{x}$ of each animal for model fitting and projections. We assumed animal weight of 5Kg.

| **Group** | **ID** | $\boldsymbol{\%}\boldsymbol{f}_{\boldsymbol{p}}$ | $\boldsymbol{D}$  **(HSPCs/Kg)** | $\boldsymbol{t}_{\boldsymbol{x}}$  **(weeks after challenge)** |
| --- | --- | --- | --- | --- |
| Control | Z09087 | 0 | 0 | N/A |
|  | Z09106 | 0 | 0 | N/A |
|  | Z09192 | 0 | 0 | N/A |
|  | Z09204 | 0 | 0 | N/A |
|  | A11201 | 0 | 0 | N/A |
| Wild-type transplantation | Z09144 | 0 | 6.45e6 | 55 |
|  | Z08214 | 0 | 4.14e6 | 58 |
|  | A11200 | 0 | 2.08e6 | 56 |
|  | Z09196 | 0 | 2.39e6 | 57 |
|  | Z09125 | 0 | 6.28e6 | 61 |
| ΔCCR5 transplantation | A11219 | 33 | 1.06e7 | 56 |
|  | T10187 | 27 | 6e6 | 57 |
|  | R10159 | 56 | 6.7e6 | 59 |
|  | T10173 | 30 | 1.48e7 | 59 |
|  | Z11151 | 60 | 1.2e7 | 63 |
|  | Z12420 | 54 | 4.82e6 | 63 |
|  | R10155 | 43 | 6e6 | 54 |
|  | Z12216 | 56 | 6.2e6 | 59 |
|  | Z12037 | 30 | 8.25e6 | 58 |
|  | Z12351 | 51 | 2.3e6 | 59 |
|  | Z13133 | 48 | 6e6 | 55 |
|  | Z12417 | 44 | 6e6 | 57 |

**Table S2.** Competing models for fitting T cell reconstitution with respective AIC values. Best fit in bold-red (lowest AIC). The AIC values presented for each statistical assumption is the lowest of 10 runs of the SAEM algorithm with different randomly selected initial guesses.

| **Model** | **Mechanistic Assumptions** | **Statistical Assumptions** | **AIC** |
| --- | --- | --- | --- |
| 1 | - Full model as in **eq. 2** in main text. | - All random effects>0. - No correlations among parameters. | -257.8 |
|  |  | - All random effects>0. - $corr(\hat{r}_{s}$,$\hat{r}_{e})\neq0.$ - $corr(N_{0}$,$E_{0})\neq0$. | -275.3 |
|  |  | - All random effects>0. - $corr(\hat{r}_{s}$,$\hat{r}_{e})\neq0.$ - $corr(N_{0}$,$E_{0})\neq0$. - $corr(K_{p}$,$E_{0})\neq0$. - $corr(N_{0}$,$K_{p})\neq0$. | -273.5 |
| 2 | - CD4^+^CCR5^+^ T cells do not downregulate CCR5 ($\lambda_{s}=0$). | - All random effects>0. - No correlations among parameters. | -221.9 |
| 3 | - Thymic export rate of naïve CD4^+^ and CD8^+^ T cells is the same ($\lambda_{e}=\lambda_{f}$). | - All random effects>0. - No correlations among parameters. | -258.5 |
|  |  | - All random effects>0. - $corr(\hat{r}_{s}$,$\hat{r}_{e})\neq0.$ - $corr(N_{0}$,$E_{0})\neq0$. | -274.3 |
|  |  | - All random effects>0. - $corr(\hat{r}_{s}$,$\hat{r}_{e})\neq0.$ - $corr(N_{0}$,$E_{0})\neq0$. - $corr(K_{p}$,$E_{0})\neq0$. - $corr(N_{0}$,$K_{p})\neq0$. | -272.2 |
| 4 | - CD4^+^CCR5^+^ T cells do not downregulate CCR5 ($\lambda_{s}=0$). - Thymic export rate of naïve CD4^+^ and CD8^+^ T cells is the same ($\lambda_{e}=\lambda_{f}$) . | - All random effects>0. - No correlations among parameters. | -231.1 |
| 5 | - CD4^+^CCR5^-^ T cells are long-lived and do not proliferate ($r_{n}=d_{n}=0$). | - All random effects>0. - No correlations among parameters. | -254.3 |
| 6 | - CD4^+^CCR5^-^ T cells are long-lived and do not proliferate ($r_{n}=d_{n}=0$). - CD4^+^CCR5^+^ T cells do not downregulate CCR5 ($\lambda_{s}=0$). | - All random effects>0. - No correlations among parameters. | -230.2 |
| 7 | - CD4^+^CCR5^-^ T cells are long-lived and do not proliferate ($r_{n}=d_{n}=0$). - Thymic export rate of naïve CD4^+^ and CD8^+^ T cells is the same ($\lambda_{e}=\lambda_{f}$). | - All random effects>0. - No correlations among parameters. | -246.7 |
| 8 | - CD4^+^CCR5^-^ T cells are long-lived and do not proliferate ($r_{n}=d_{n}=0$). - Thymic export rate of naïve CD4^+^ and CD8^+^ T cells is the same ($\lambda_{e}=\lambda_{f}$). - CD4^+^CCR5^+^ T cells do not downregulate CCR5 ($\lambda_{s}=0$). | - All random effects>0. - No correlations among parameters. | -209.1 |
| 9 | - CD4^+^CCR5^-^ T cells do not proliferate ($r_{n}=0$). | - All random effects>0. - No correlations among parameters. | -255.7 |
|  |  | - All random effects>0. - $corr(\hat{r}_{s}$,$\hat{r}_{e})\neq0.$ - $corr(N_{0}$,$E_{0})\neq0$. | -277 |
|  |  | - All random effects>0. - $corr(\hat{r}_{s}$,$\hat{r}_{e})\neq0.$ - $corr(N_{0}$,$E_{0})\neq0$. - $corr(K_{p}$,$E_{0})\neq0$. - $corr(N_{0}$,$K_{p})\neq0$. | -273 |
| 10 | - CD4^+^CCR5^-^ T cells do not proliferate ($r_{n}=0$). - CD4^+^CCR5^+^ T cells do not downregulate CCR5 ($\lambda_{s}=0$). | - All random effects>0. - No correlations among parameters. | -233.3 |
| 11 | - CD4^+^CCR5^-^ T cells do not proliferate ($r_{n}=0$). - Thymic export rate of naïve CD4^+^ and CD8^+^ T cells is the same ($\lambda_{e}=\lambda_{f}$). | - All random effects>0, - no correlations. | -254.5 |
|  |  | - All random effects>0. - $corr(\hat{r}_{s}$,$\hat{r}_{e})\neq0.$ - $corr(N_{0}$,$E_{0})\neq0$. | -279.1 |
|  |  | - All random effects>0. - $corr(\hat{r}_{s}$,$\hat{r}_{e})\neq0.$ - $corr(N_{0}$,$E_{0})\neq0$. - $corr(K_{p}$,$E_{0})\neq0$. - $corr(N_{0}$,$K_{p})\neq0$. | -285.3 |
|  |  | - Random effects equal to zero for parameters: $\hat{r}_{m}$, $\lambda_{n}$, $\lambda_{s}$, $\hat{d}_{n}$, $K_{s}$, and $K_{e}.$ - All random effects>0. - $corr(\hat{r}_{s}$,$\hat{r}_{e})\neq0.$ - $corr(N_{0}$,$E_{0})\neq0$. - $corr(K_{p}$,$E_{0})\neq0$. - $corr(N_{0}$,$K_{p})\neq0$. | -297.4 |
|  |  | - Random effects equal to zero for parameters: $\hat{r}_{m}$, $\lambda_{e}$, $\hat{d}_{n}$, $K_{s}$, and $K_{e}.$ - $corr(\hat{r}_{s}$,$\hat{r}_{e})\neq0.$ - $corr(N_{0}$,$E_{0})\neq0$. - $corr(K_{p}$,$E_{0})\neq0$. - $corr(N_{0}$,$K_{p})\neq0$. | **-300.3** |
| 12 | - CD4^+^CCR5^-^ T cells do not proliferate ($r_{n}=0$). - Thymic export rate of naïve CD4^+^ and CD8^+^ T cells is the same ($\lambda_{e}=\lambda_{f}$). - CD4^+^CCR5^+^ T cells do not downregulate CCR5 ($\lambda_{s}=0$). | - All random effects>0. - No correlations among parameters. | -236.5 |

**Table S3.** Population parameter estimates for the best fits of the model in **eq. 2** in the main text (lowest AIC in **Table S2**) to the T cell reconstitution dynamics. RSE: relative standard error. Empty fields represent a standard deviation of random effects, $\sigma_{\psi}$, fixed to zero. Values of $\bar{\psi}$ for $K_{p},N\left( t_{0} \right),S\left( t_{0} \right),M\left( t_{0} \right),$ and $E\left( t_{0} \right)$ shown here are in log_10_ cell counts/ μL assuming a blood volume of of 3×10^5^ μL (calculated assuming blood:weight ratio of 60mL/Kg and body weight of 5Kg). Red values represent an RSE greater than 100% implying that the number of data points may not be enough to estimate the respective parameter.

| **Parameter** | $\bar{\boldsymbol{\psi}}$ | $\boldsymbol{\sigma}_{\boldsymbol{\psi}}$ | **%RSE for:** | |
| --- | --- | --- | --- | --- |
|  |  |  | $\bar{\boldsymbol{\psi}}$ | $\boldsymbol{\sigma}_{\boldsymbol{\psi}}$ |
| ${\hat{\boldsymbol{r}}}_{\boldsymbol{p}}$ | 0.05 | 0.39 | 20 | 23 |
| ${\hat{\boldsymbol{r}}}_{\boldsymbol{s}}$ | 0.11 | 0.41 | 10 | 18 |
| ${\hat{\boldsymbol{r}}}_{\boldsymbol{m}}$ | 0.03 |  | 51 |  |
| ${\hat{\boldsymbol{r}}}_{\boldsymbol{e}}$ | 0.08 | 0.56 | 15 | 18 |
| ${\hat{\boldsymbol{d}}}_{\boldsymbol{n}}$ | 8.2 |  | 36 |  |
| $\boldsymbol{\lambda}_{\boldsymbol{e}}\boldsymbol{=}\boldsymbol{\lambda}_{\boldsymbol{f}}$ | 0.003 |  | 43 |  |
| $\boldsymbol{\lambda}_{\boldsymbol{s}}$ | 0.014 | 0.2 | 24 | 130 |
| $\boldsymbol{\lambda}_{\boldsymbol{n}}$ | 0.003 | 0.27 | 27 | 40 |
| $\boldsymbol{\lambda}_{\boldsymbol{m}}$ | 0.07 |  | 27 |  |
| $\boldsymbol{K}_{\boldsymbol{p}}$ | 3.2 | 0.23 | 1 | 16 |
| $\boldsymbol{K}_{\boldsymbol{s}}$ | 0.13 |  | 21 |  |
| $\boldsymbol{K}_{\boldsymbol{m}}$ | 0.75 | 0.28 | 28 | 22 |
| $\boldsymbol{K}_{\boldsymbol{e}}$ | 0.15 |  | 19 |  |
| $\boldsymbol{N(}\boldsymbol{t}_{\boldsymbol{0}}\boldsymbol{)}$ | 1.9 | 0.13 | 1 | 27 |
| $\boldsymbol{S(}\boldsymbol{t}_{\boldsymbol{0}}\boldsymbol{)}$ | 0.64 | 0.27 | 1 | 23 |
| $\boldsymbol{M(}\boldsymbol{t}_{\boldsymbol{0}}\boldsymbol{)}$ | 1.0 | 0.14 | 2 | 73 |
| $\boldsymbol{E(}\boldsymbol{t}_{\boldsymbol{0}}\boldsymbol{)}$ | 1.3 | 0.4 | 2 | 21 |
|  | **Parameter value** | | **%RSE** | |
| $\boldsymbol{corr(}{\hat{\boldsymbol{r}}}_{\boldsymbol{s}}$**,**${\hat{\boldsymbol{r}}}_{\boldsymbol{e}}\boldsymbol{)}$ | 0.87 | | 9 | |
| $\boldsymbol{corr}\boldsymbol{(}\boldsymbol{N}_{\boldsymbol{0}}$**,**$\boldsymbol{E}_{\boldsymbol{0}}\boldsymbol{)}$ | 0.99 | | 14 | |
| $\boldsymbol{corr}\boldsymbol{(}\boldsymbol{K}_{\boldsymbol{p}}$**,**$\boldsymbol{E}_{\boldsymbol{0}}\boldsymbol{)}$ | 0.8 | | 15.5 | |
| $\boldsymbol{corr}\boldsymbol{(}\boldsymbol{N}_{\boldsymbol{0}}$**,**$\boldsymbol{K}_{\boldsymbol{p}}\boldsymbol{)}$ | 0.74 | | 26 | |
| $\boldsymbol{\sigma}_{\boldsymbol{N}}$ | 0.2 | | 4 | |
| $\boldsymbol{\sigma}_{\boldsymbol{S}}$ | 0.16 | | 4 | |
| $\boldsymbol{\sigma}_{\boldsymbol{C}}$ | 0.19 | | 4 | |
| $\boldsymbol{\sigma}_{\boldsymbol{E}}$ | 0.18 | | 11 | |
| $\boldsymbol{\sigma}_{\boldsymbol{M}}$ | 0.21 | | 12 | |

**Table S4.** Individual parameter estimates for the best fits of the model in **eq. 2** in the main text (lowest AIC in **Table S2**) to the T cell reconstitution dynamics. Values obtained for $N\left( t_{0} \right),S\left( t_{0} \right),M\left( t_{0} \right),$ and $E\left( t_{0} \right)$ shown here are in log_10_ cell counts/ μL assuming a blood volume of of 3×10^5^ μL (calculated assuming blood:weight ratio of 60mL/Kg and body weight of 5Kg). Initial values for the control group where obtained assuming steady state.

|  | **Control** | | | | | **WT-Transplant** | | | | | **ΔCCR5-Transplant** | | | | | | | | | | | |
| --- | --- | --- | --- | --- | --- | --- | --- | --- | --- | --- | --- | --- | --- | --- | --- | --- | --- | --- | --- | --- | --- | --- |
| **Par.**  **ID** | **Z09087** | **Z09106** | **Z09192** | **Z09204** | **A11201** | **Z09144** | **Z08214** | **A11200** | **Z09196** | **Z09125** | **A11219** | **T10187** | **R10159** | **T10173** | **Z11151** | **Z12420** | **R10155** | **Z12216** | **Z12037** | **Z12351** | **Z13133** | **Z12417** |
| ${\hat{\boldsymbol{r}}}_{\boldsymbol{p}}^{\boldsymbol{j}}$  **(1/day)** | 0.05 | 0.05 | 0.05 | 0.05 | 0.05 | 0.06 | 0.04 | 0.04 | 0.07 | 0.07 | 0.03 | 0.05 | 0.04 | 0.04 | 0.05 | 0.08 | 0.05 | 0.09 | 0.05 | 0.05 | 0.04 | 0.08 |
| ${\hat{\boldsymbol{r}}}_{\boldsymbol{s}}^{\boldsymbol{j}}$  **(1/day)** | 0.10 | 0.10 | 0.10 | 0.07 | 0.11 | 0.12 | 0.14 | 0.07 | 0.20 | 0.07 | 0.12 | 0.12 | 0.14 | 0.19 | 0.07 | 0.09 | 0.09 | 0.08 | 0.22 | 0.06 | 0.14 | 0.14 |
| ${\hat{\boldsymbol{r}}}_{\boldsymbol{m}}^{\boldsymbol{j}}$ **(1/day)** | 0.03 | 0.03 | 0.03 | 0.03 | 0.03 | 0.03 | 0.03 | 0.03 | 0.03 | 0.03 | 0.03 | 0.03 | 0.03 | 0.03 | 0.03 | 0.03 | 0.03 | 0.03 | 0.03 | 0.03 | 0.03 | 0.03 |
| ${\hat{\boldsymbol{r}}}_{\boldsymbol{e}}^{\boldsymbol{j}}$  **(1/day)** | 0.05 | 0.07 | 0.07 | 0.04 | 0.07 | 0.10 | 0.10 | 0.06 | 0.19 | 0.04 | 0.07 | 0.11 | 0.11 | 0.15 | 0.05 | 0.10 | 0.05 | 0.05 | 0.17 | 0.03 | 0.13 | 0.15 |
| ${\hat{\boldsymbol{d}}}_{\boldsymbol{n}}^{\boldsymbol{j}}$ **(1/day)** | 0.02 | 0.04 | 0.03 | 0.03 | 0.03 | 0.02 | 0.03 | 0.03 | 0.02 | 0.02 | 0.04 | 0.03 | 0.03 | 0.03 | 0.03 | 0.03 | 0.03 | 0.02 | 0.03 | 0.03 | 0.04 | 0.03 |
| $\boldsymbol{\lambda}_{\boldsymbol{e}}^{\boldsymbol{j}}$  **(1/day)** | 0.003 | 0.003 | 0.003 | 0.003 | 0.003 | 0.003 | 0.003 | 0.003 | 0.003 | 0.003 | 0.003 | 0.003 | 0.003 | 0.003 | 0.003 | 0.003 | 0.003 | 0.003 | 0.003 | 0.003 | 0.003 | 0.003 |
| $\boldsymbol{\lambda}_{\boldsymbol{n}}^{\boldsymbol{j}}$  **(1/day)** | 0.002 | 0.004 | 0.003 | 0.003 | 0.003 | 0.003 | 0.003 | 0.003 | 0.002 | 0.003 | 0.004 | 0.003 | 0.003 | 0.003 | 0.003 | 0.004 | 0.004 | 0.002 | 0.003 | 0.003 | 0.004 | 0.003 |
| $\boldsymbol{\lambda}_{\boldsymbol{s}}^{\boldsymbol{j}}$  **(1/day)** | 0.01 | 0.01 | 0.01 | 0.01 | 0.01 | 0.02 | 0.01 | 0.01 | 0.02 | 0.01 | 0.01 | 0.01 | 0.01 | 0.01 | 0.01 | 0.01 | 0.02 | 0.01 | 0.02 | 0.01 | 0.01 | 0.01 |
| $\boldsymbol{\lambda}_{\boldsymbol{m}}^{\boldsymbol{j}}$ **(1/day)** | 0.07 | 0.07 | 0.07 | 0.07 | 0.07 | 0.07 | 0.07 | 0.07 | 0.07 | 0.07 | 0.07 | 0.07 | 0.07 | 0.07 | 0.07 | 0.07 | 0.07 | 0.07 | 0.07 | 0.07 | 0.07 | 0.07 |
| $\boldsymbol{K}_{\boldsymbol{p}}^{\boldsymbol{j}}$  $\left( \boldsymbol{cells} \right)$ | 10^8.9^ | 10^8.8^ | 10^9.0^ | 10^8.8^ | 10^8.7^ | 10^8.6^ | 10^8.7^ | 10^8.7^ | 10^8.5^ | 10^8.7^ | 10^8.5^ | 10^8.7^ | 10^8.8^ | 10^8.5^ | 10^8.8^ | 10^8.5^ | 10^8.9^ | 10^8.8^ | 10^8.0^ | 10^8.4^ | 10^8.2^ | 10^8.7^ |
| $\boldsymbol{K}_{\boldsymbol{s}}^{\boldsymbol{j}}$ $\left( \frac{\boldsymbol{cells}}{\boldsymbol{\mu L}} \right)$ | 2134 | 1535 | 2350 | 1625 | 1274 | 923 | 1214 | 1173 | 739 | 1359 | 735 | 1117 | 1425 | 766 | 1636 | 779 | 1790 | 1654 | 275 | 650 | 407 | 1124 |
| $\boldsymbol{K}_{\boldsymbol{m}}^{\boldsymbol{j}}$ $\left( \frac{\boldsymbol{cells}}{\boldsymbol{\mu L}} \right)$ | 678 | 718 | 862 | 277 | 439 | 56 | 398 | 315 | 233 | 307 | 112 | 225 | 327 | 113 | 289 | 154 | 404 | 367 | 52 | 126 | 112 | 281 |
| $\boldsymbol{K}_{\boldsymbol{e}}^{\boldsymbol{j}}$ $\left( \frac{\boldsymbol{cells}}{\boldsymbol{\mu L}} \right)$ | 2033 | 1462 | 2239 | 1548 | 1214 | 880 | 1156 | 1117 | 704 | 1294 | 700 | 1065 | 1358 | 730 | 1559 | 742 | 1706 | 1576 | 262 | 620 | 388 | 1071 |
| $\boldsymbol{N}^{\boldsymbol{j}}\left( \boldsymbol{t}_{\boldsymbol{0}} \right) \left( \frac{\boldsymbol{cells}}{\boldsymbol{\mu L}} \right)$ | 1452 | 621 | 1350 | 1253 | 832 | 65.6 | 58.6 | 94.6 | 54.5 | 83.4 | 58.2 | 76.3 | 80.1 | 80.0 | 89.0 | 53.4 | 81.8 | 116.9 | 33.9 | 54.8 | 56.8 | 61.9 |
| $\boldsymbol{S}^{\boldsymbol{j}}\left( \boldsymbol{t}_{\boldsymbol{0}} \right) \left( \frac{\boldsymbol{cells}}{\boldsymbol{\mu L}} \right)$ | 109 | 82 | 135 | 164 | 69 | 2.7 | 1.9 | 8.3 | 1.6 | 2.7 | 3.5 | 6.8 | 7.4 | 8.5 | 6.3 | 4.5 | 3.6 | 10.2 | 2.2 | 6.3 | 4.3 | 3.5 |
| $\boldsymbol{M}^{\boldsymbol{j}}\boldsymbol{(}\boldsymbol{t}_{\boldsymbol{0}}\boldsymbol{)}$ $\left( \frac{\boldsymbol{cells}}{\boldsymbol{\mu L}} \right)$ | 320 | 412 | 496 | 155 | 258 | 8.9 | 9.6 | 14.4 | 11.1 | 10.4 | 11.0 | 10.6 | 10.8 | 10.9 | 11.8 | 11.9 | 11.0 | 10.7 | 8.8 | 9.2 | 13.8 | 9.1 |
| $\boldsymbol{E}^{\boldsymbol{j}}\boldsymbol{(}\boldsymbol{t}_{\boldsymbol{0}}\boldsymbol{)}$ $\left( \frac{\boldsymbol{cells}}{\boldsymbol{\mu L}} \right)$ | 1000 | 957 | 1191 | 621 | 561 | 16.3 | 12.7 | 48.4 | 9.1 | 35.3 | 11.0 | 26.1 | 31.9 | 27.1 | 44.3 | 8.7 | 35.8 | 95.6 | 1.9 | 9.0 | 8.8 | 14.5 |

**Table S5.** Competing models for fitting T cell and viral dynamics (**eqs. 2-3** in main text) using the best model in **Table S2** and fixing parameter values as in **Table S3**, with AIC values. Best fit in bold-red (lowest AIC).

| **Model** | **Mechanistic Assumptions** | **Statistical Assumptions** | **AIC** |
| --- | --- | --- | --- |
| 1 | - SHIV-specific CD8^+^ T cells reduce virus production only ($\theta>0, \kappa=0$). - Immunity is lost during TBI: ($\omega_{8}$, $I_{50}$ and $d_{h}$ different during acute infection and after ATI). - SHIV-infection enhances activation of CD4^+^CCR5^-^ T cells leading to replenishment of CD4^+^CCR5^+^ T cells, and transient reduction of the CD4^+^CCR5^-^ compartment after ATI ($\omega_{4}>0$). | - $\sigma_{t_{sa}}=1, \sigma_{\pi}=0.5$. - ${\psi_{j}^{ATI}=10}^{{\bar{\psi}+n}_{j}+\varsigma_{\psi,\mathrm{ATI}}}$ for $\omega_{8}$and $I_{50}$, and $\psi_{j}^{ATI}=\bar{\psi}e^{n_{j}+\varsigma_{\psi,\mathrm{ATI}}}$ for $d_{h}$. - $\theta=1/\mu L$. - $corr(I_{50}$,$d_{h})\neq0.$ - $corr(\omega_{8}$,$d_{h})\neq0$. - $corr(I_{50},\omega_{8})\neq0$. - $corr(\pi$,$\beta)\neq0$. | -1697.5 |
|  |  | - $\sigma_{t_{sa}}=1, \sigma_{\pi}=0.5$. - ${\psi_{j}^{ATI}=10}^{{\bar{\psi}+n}_{j}+\varsigma_{\psi,\mathrm{ATI}}}$ for $\omega_{8}$and $I_{50}$, and $\psi_{j}^{ATI}=\bar{\psi}e^{n_{j}+\varsigma_{\psi,\mathrm{ATI}}}$ for $d_{h}$. - $\theta=1/\mu L$. - $corr(\hat{r}_{s},\lambda_{n})\neq0.$ - $corr(\hat{r}_{e},\lambda_{n})\neq0.$ - $corr(I_{50}$,$d_{h})\neq0.$ - $corr(\omega_{8}$,$d_{h})\neq0$. - $corr(I_{50},\omega_{8})\neq0$. - $corr(\pi$,$\beta)\neq0$. - $corr(k_{t}$,$k_{h})\neq0.$ - $corr(\pi$,$d_{h})\neq0$. - $corr(I_{50},\beta)\neq0$. - $corr(\pi$,$I_{50})\neq0$. - $corr(\omega_{8}$,$\beta)\neq0$. - $corr(\pi$,$\omega_{8})\neq0$. - $t_{sa}^{j}$ for ΔCCR5 and transplant groups was modeled as $\psi_{j}=\bar{\psi}e^{n_{j}+\varsigma_{t_{sa},\Delta CCR5}}$ and $\psi_{j}=\bar{\psi}e^{n_{j}+\varsigma_{t_{sa},\mathrm{WT}}}$, respectively. - $K_{p}^{j}$ for ΔCCR5 and transplant groups was modeled as $\psi_{j}={10}^{{\bar{\psi}+n}_{j}+\varsigma_{K_{p},\Delta CCR5}}$ and $\psi_{j}={10}^{{\bar{\psi}+n}_{j}+\varsigma_{K_{p},WT}}$ respectively. | -1725.8 |
|  |  | - $\sigma_{t_{sa}}=1, \sigma_{\pi}=0.5$. - ${\psi_{j}^{ATI}=10}^{{\bar{\psi}+n}_{j}+\varsigma_{\psi,\mathrm{ATI}}}$ for $\omega_{8}$and $I_{50}$, and $\psi_{j}^{ATI}=\bar{\psi}e^{n_{j}+\varsigma_{\psi,\mathrm{ATI}}}$ for $d_{h}$. - $\theta=1/\mu L$. - $corr(\hat{r}_{s},\lambda_{n})\neq0.$ - $corr(\hat{r}_{e},\lambda_{n})\neq0.$ - $corr(\hat{r}_{s},\beta)\neq0.$ - $corr(\hat{r}_{e},\beta)\neq0.$ - $corr(\hat{r}_{s},\pi)\neq0.$ - $corr(\hat{r}_{e},\pi)\neq0.$ - $corr(\beta,\lambda_{n})\neq0.$ - $corr(\pi,\lambda_{n})\neq0.$ - $corr(I_{50}$,$d_{h})\neq0.$ - $corr(\omega_{8}$,$d_{h})\neq0$. - $corr(I_{50},\omega_{8})\neq0$. - $corr(\pi$,$\beta)\neq0$. - $corr(k_{t}$,$k_{h})\neq0.$ - $t_{sa}^{j}$ for ΔCCR5 and transplant groups was modeled as $\psi_{j}=\bar{\psi}e^{n_{j}+\varsigma_{t_{sa},\Delta CCR5}}$ and $\psi_{j}=\bar{\psi}e^{n_{j}+\varsigma_{t_{sa},\mathrm{WT}}}$, respectively. - $K_{p}^{j}$ for ΔCCR5 and transplant groups was modeled as $\psi_{j}={10}^{{\bar{\psi}+n}_{j}+\varsigma_{K_{p},\Delta CCR5}}$ and $\psi_{j}={10}^{{\bar{\psi}+n}_{j}+\varsigma_{K_{p},WT}}$ respectively. | -1722.7 |
|  |  | - $\sigma_{t_{sa}}=1, \sigma_{\pi}=0.5$. - ${\psi_{j}^{ATI}=10}^{{\bar{\psi}+n}_{j}+\varsigma_{\psi,\mathrm{ATI}}}$ for $\omega_{8}$and $I_{50}$, and $\psi_{j}^{ATI}=\bar{\psi}e^{n_{j}+\varsigma_{\psi,\mathrm{ATI}}}$ for $d_{h}$. - $\theta=1/\mu L$. - $corr(\hat{r}_{s},\lambda_{n})\neq0.$ - $corr(\hat{r}_{e},\lambda_{n})\neq0.$ - $corr(I_{50}$,$d_{h})\neq0.$ - $corr(\omega_{8}$,$d_{h})\neq0$. - $corr(I_{50},\omega_{8})\neq0$. - $corr(\pi$,$\beta)\neq0$. - $corr(k_{t}$,$k_{h})\neq0.$ - $t_{sa}^{j}$ for ΔCCR5 and transplant groups was modeled as $\psi_{j}=\bar{\psi}e^{n_{j}+\varsigma_{t_{sa},\Delta CCR5}}$ and $\psi_{j}=\bar{\psi}e^{n_{j}+\varsigma_{t_{sa},\mathrm{WT}}}$, respectively. - $K_{p}^{j}$ for ΔCCR5 and transplant groups was modeled as $\psi_{j}={10}^{{\bar{\psi}+n}_{j}+\varsigma_{K_{p},\Delta CCR5}}$ and $\psi_{j}={10}^{{\bar{\psi}+n}_{j}+\varsigma_{K_{p},WT}}$ respectively. | **-1774.9** |
| 2 | - SHIV-specific CD8^+^ T cells kill SHIV-infected cells only ($\theta=0, \kappa>0$). - Immunity is lost during TBI: ($\omega_{8}$, $I_{50}$ and $d_{h}$ different during acute infection and after ATI). - SHIV-infection enhances activation of CD4^+^CCR5^-^ T cells leading to replenishment of CD4^+^CCR5^+^ T cells, and transient reduction of the CD4^+^CCR5^-^ compartment after ATI ($\omega_{4}>0$). | - $\sigma_{t_{sa}}=1, \sigma_{\pi}=0.5$. - ${\psi_{j}^{ATI}=10}^{{\bar{\psi}+n}_{j}+\varsigma_{\psi,\mathrm{ATI}}}$ for $\omega_{8}$and $I_{50}$, and $\psi_{j}^{ATI}=\bar{\psi}e^{n_{j}+\varsigma_{\psi,\mathrm{ATI}}}$ for $d_{h}$. - $\kappa=1/\mu L$. - $corr(\hat{r}_{s},\lambda_{n})\neq0.$ - $corr(\hat{r}_{e},\lambda_{n})\neq0.$ - $corr(\hat{r}_{s},\beta)\neq0.$ - $corr(\hat{r}_{e},\beta)\neq0.$ - $corr(\hat{r}_{s},\pi)\neq0.$ - $corr(\hat{r}_{e},\pi)\neq0.$ - $corr(\beta,\lambda_{n})\neq0.$ - $corr(\pi,\lambda_{n})\neq0.$ - $corr(I_{50}$,$d_{h})\neq0.$ - $corr(\omega_{8}$,$d_{h})\neq0$. - $corr(I_{50},\omega_{8})\neq0$. - $corr(\pi$,$\beta)\neq0$. - $corr(k_{t}$,$k_{h})\neq0.$ - $t_{sa}^{j}$ for ΔCCR5 and transplant groups was modeled as $\psi_{j}=\bar{\psi}e^{n_{j}+\varsigma_{t_{sa},\Delta CCR5}}$ and $\psi_{j}=\bar{\psi}e^{n_{j}+\varsigma_{t_{sa},\mathrm{WT}}}$, respectively. - $K_{p}^{j}$ for ΔCCR5 and transplant groups was modeled as $\psi_{j}={10}^{{\bar{\psi}+n}_{j}+\varsigma_{K_{p},\Delta CCR5}}$ and $\psi_{j}={10}^{{\bar{\psi}+n}_{j}+\varsigma_{K_{p},WT}}$ respectively. | -436.1 |
|  |  | - $\sigma_{t_{sa}}=1, \sigma_{\pi}=0.5$. - ${\psi_{j}^{ATI}=10}^{{\bar{\psi}+n}_{j}+\varsigma_{\psi,\mathrm{ATI}}}$ for $\omega_{8}$and $I_{50}$, and $\psi_{j}^{ATI}=\bar{\psi}e^{n_{j}+\varsigma_{\psi,\mathrm{ATI}}}$ for $d_{h}$. - $\kappa=1$ - $corr(\hat{r}_{s},\lambda_{n})\neq0.$ - $corr(\hat{r}_{e},\lambda_{n})\neq0.$ - $corr(I_{50}$,$d_{h})\neq0.$ - $corr(\omega_{8}$,$d_{h})\neq0$. - $corr(I_{50},\omega_{8})\neq0$. - $corr(\pi$,$\beta)\neq0$. - $corr(k_{t}$,$k_{h})\neq0.$ - $t_{sa}^{j}$ for ΔCCR5 and transplant groups was modeled as $\psi_{j}=\bar{\psi}e^{n_{j}+\varsigma_{t_{sa},\Delta CCR5}}$ and $\psi_{j}=\bar{\psi}e^{n_{j}+\varsigma_{t_{sa},\mathrm{WT}}}$, respectively. - $K_{p}^{j}$ for ΔCCR5 and transplant groups was modeled as $\psi_{j}={10}^{{\bar{\psi}+n}_{j}+\varsigma_{K_{p},\Delta CCR5}}$ and $\psi_{j}={10}^{{\bar{\psi}+n}_{j}+\varsigma_{K_{p},WT}}$ respectively. | -1511.5 |
|  |  | - $\sigma_{t_{sa}}=1, \sigma_{\pi}=0.5$. - ${\psi_{j}^{ATI}=10}^{{\bar{\psi}+n}_{j}+\varsigma_{\psi,\mathrm{ATI}}}$ for $\omega_{8}$and $I_{50}$, and $\psi_{j}^{ATI}=\bar{\psi}e^{n_{j}+\varsigma_{\psi,\mathrm{ATI}}}$ for $d_{h}$. - $\kappa$ modeled as ${\psi_{j}=10}^{{\bar{\psi}+n}_{j}}$ - $corr(\hat{r}_{s},\lambda_{n})\neq0.$ - $corr(\hat{r}_{e},\lambda_{n})\neq0.$ - $corr(I_{50}$,$d_{h})\neq0.$ - $corr(\omega_{8}$,$d_{h})\neq0$. - $corr(I_{50},\omega_{8})\neq0$. - $corr(\pi$,$\beta)\neq0$. - $corr(k_{t}$,$k_{h})\neq0.$ - $t_{sa}^{j}$ for ΔCCR5 and transplant groups was modeled as $\psi_{j}=\bar{\psi}e^{n_{j}+\varsigma_{t_{sa},\Delta CCR5}}$ and $\psi_{j}=\bar{\psi}e^{n_{j}+\varsigma_{t_{sa},\mathrm{WT}}}$, respectively. - $K_{p}^{j}$ for ΔCCR5 and transplant groups was modeled as $\psi_{j}={10}^{{\bar{\psi}+n}_{j}+\varsigma_{K_{p},\Delta CCR5}}$ and $\psi_{j}={10}^{{\bar{\psi}+n}_{j}+\varsigma_{K_{p},WT}}$ respectively. | -1630.8 |
| 3 | - SHIV-specific CD8^+^ T cells reduce virus production only ($\theta>0, \kappa=0$). - Immunity is *not* lost during TBI ($\omega_{8},I_{50},d_{h}$ equal during acute infection and after ATI). - SHIV-infection enhances activation of CD4^+^CCR5^-^ T cells leading to replenishment of CD4^+^CCR5^+^ T cells, and transient reduction of the CD4^+^CCR5^-^ compartment after ATI ($\omega_{4}>0$). | - $\sigma_{t_{sa}}=1, \sigma_{\pi}=0.5$. - $\theta=1/\mu L$. - $corr(\hat{r}_{s},\lambda_{n})\neq0.$ - $corr(\hat{r}_{e},\lambda_{n})\neq0.$ - $corr(I_{50}$,$d_{h})\neq0.$ - $corr(\omega_{8}$,$d_{h})\neq0$. - $corr(I_{50},\omega_{8})\neq0$. - $corr(\pi$,$\beta)\neq0$. - $corr(k_{t}$,$k_{h})\neq0.$ - $t_{sa}^{j}$ for ΔCCR5 and transplant groups was modeled as $\psi_{j}=\bar{\psi}e^{n_{j}+\varsigma_{t_{sa},\Delta CCR5}}$ and $\psi_{j}=\bar{\psi}e^{n_{j}+\varsigma_{t_{sa},\mathrm{WT}}}$, respectively. - $K_{p}^{j}$ for ΔCCR5 and transplant groups was modeled as $\psi_{j}={10}^{{\bar{\psi}+n}_{j}+\varsigma_{K_{p},\Delta CCR5}}$ and $\psi_{j}={10}^{{\bar{\psi}+n}_{j}+\varsigma_{K_{p},WT}}$ respectively. | -1177.2 |
| 4 | - SHIV-specific CD8^+^ T cells reduce virus production only ($\theta>0, \kappa=0$). - Immunity is lost during TBI: ($\omega_{8}$, $I_{50}$ and $d_{h}$ different during acute infection and after ATI). - SHIV-infection does not enhance activation of CD4^+^CCR5^-^ T cells or replenishment of CD4^+^CCR5^+^ T cells ($\omega_{4}=0$). | - $\sigma_{t_{sa}}=1, \sigma_{\pi}=0.5$. - ${\psi_{j}^{ATI}=10}^{{\bar{\psi}+n}_{j}+\varsigma_{\psi,\mathrm{ATI}}}$ for $\omega_{8}$and $I_{50}$, and $\psi_{j}^{ATI}=\bar{\psi}e^{n_{j}+\varsigma_{\psi,\mathrm{ATI}}}$ for $d_{h}$. - $\theta=1/\mu L$. - $corr(\hat{r}_{s},\lambda_{n})\neq0.$ - $corr(\hat{r}_{e},\lambda_{n})\neq0.$ - $corr(I_{50}$,$d_{h})\neq0.$ - $corr(\omega_{8}$,$d_{h})\neq0$. - $corr(I_{50},\omega_{8})\neq0$. - $corr(\pi$,$\beta)\neq0$. - $corr(k_{t}$,$k_{h})\neq0.$ - $t_{sa}^{j}$ for ΔCCR5 and transplant groups was modeled as $\psi_{j}=\bar{\psi}e^{n_{j}+\varsigma_{t_{sa},\Delta CCR5}}$ and $\psi_{j}=\bar{\psi}e^{n_{j}+\varsigma_{t_{sa},\mathrm{WT}}}$, respectively. - $K_{p}^{j}$ for ΔCCR5 and transplant groups was modeled as $\psi_{j}={10}^{{\bar{\psi}+n}_{j}+\varsigma_{K_{p},\Delta CCR5}}$ and $\psi_{j}={10}^{{\bar{\psi}+n}_{j}+\varsigma_{K_{p},WT}}$ respectively. | -1224.6 |

**Table S6.** Population parameter estimates for the fits of the model with lowest AIC in **Table S5** to the T cell and virus dynamics. RSE: relative standard error. Empty fields represent cases when the standard deviation of random effects, $\boldsymbol{\sigma}_{\boldsymbol{\psi}}$ , was fixed to zero. Values of $\bar{\psi}$ for $\beta,\omega_{4},\omega_{8}$ and $I_{50}$ shown here are transformed assuming a blood volume of 3×10^5^ μL (calculated assuming blood:weight ratio of 60mL/Kg and body weight of 5Kg). Red values represent an RSE greater than 100% implying that the number of data points may not be enough to estimate the respective parameter.

| **Parameter** | $\bar{\boldsymbol{\psi}}$ | $\boldsymbol{\sigma}_{\boldsymbol{\psi}}$ | **%RSE for:** | |
| --- | --- | --- | --- | --- |
|  |  |  | $\bar{\boldsymbol{\psi}}$ | $\boldsymbol{\sigma}_{\boldsymbol{\psi}}$ |
| $\boldsymbol{K}_{\boldsymbol{p}}$ | 3.2 | 0.17 | 1 | 25 |
|  | $\varsigma_{K_{p},\Delta CCR5}=-0.2$ |  | 49 |  |
|  | $\varsigma_{K_{p},\Delta CCR5}=-0.1$ |  | 49 |  |
| $\boldsymbol{\beta}$ | -3.8 | 0.3 | 1.9 | 14 |
| $\boldsymbol{\pi}$ | 5.2 | 0.5 | 1.9 |  |
| $\boldsymbol{\omega}_{\boldsymbol{4}}$ | -2.4 | 0.4 | 1 | 20 |
| $\boldsymbol{\omega}_{\boldsymbol{8}}$ | -3.2 | 0.7 | 1.8 | 17 |
|  | $\varsigma_{\omega_{8},\mathrm{ATI}}=1.35$ | 0.9 | 20 | 22 |
| $\boldsymbol{I}_{\mathbf{50}}$ | 0.8 | 0.7 | 2 | 19 |
|  | $\varsigma_{I_{50},\mathrm{ATI}}=0.55$ | 0.2 | 23 | 82 |
| $\boldsymbol{d}_{\boldsymbol{h}}$ | 0.003 | 0.9 | 21 | 21 |
|  | $\varsigma_{d_{h},\mathrm{ATI}}=2.9$ | 1.5 | 23 | 21 |
| $\boldsymbol{t}_{\boldsymbol{sa}}$ | 3.1 | 1 | 20 |  |
|  | $\varsigma_{t_{sa},WT}=1.3$ |  | 45 |  |
|  | $\varsigma_{t_{sa},\Delta CCR5}=1.8$ |  | 27 |  |
| $\boldsymbol{k}_{\boldsymbol{T}}$ | 0.52 | 0.1 | 4 | 20 |
| $\boldsymbol{k}_{\boldsymbol{H}}$ | 1.6 | 0.1 | 4 | 20 |
|  | **Parameter value** | | **%RSE** | |
| $\boldsymbol{corr(}{\hat{\boldsymbol{r}}}_{\boldsymbol{s}}\boldsymbol{,}\boldsymbol{\lambda}_{\boldsymbol{n}}\boldsymbol{)}$ | 0.6 | | 20 | |
| $\boldsymbol{corr(}{\hat{\boldsymbol{r}}}_{\boldsymbol{e}}\boldsymbol{,}\boldsymbol{\lambda}_{\boldsymbol{n}}\boldsymbol{)}$ | 0.8 | | 11 | |
| $\boldsymbol{corr(}\boldsymbol{I}_{\boldsymbol{50}}$**,**$\boldsymbol{d}_{\boldsymbol{h}}\boldsymbol{)}$ | -0.7 | | 27 | |
| $\boldsymbol{corr}\boldsymbol{(}\boldsymbol{\omega}_{\boldsymbol{8}}$**,**$\boldsymbol{d}_{\boldsymbol{h}}\boldsymbol{)}$ | 0.6 | | 34 | |
| $\boldsymbol{corr}\boldsymbol{(}\boldsymbol{I}_{\boldsymbol{50}}\mathbf{,}\boldsymbol{\omega}_{\boldsymbol{8}}\boldsymbol{)}$ | -0.62 | | 24 | |
| $\boldsymbol{corr}\boldsymbol{(}\boldsymbol{\pi}$**,**$\boldsymbol{\beta}\boldsymbol{)}$ | -0.89 | | 5 | |
| $\boldsymbol{corr}\boldsymbol{(}\boldsymbol{k}_{\boldsymbol{T}}$**,**$\boldsymbol{k}_{\boldsymbol{H}}\boldsymbol{)}$ | -0.93 | | 6 | |
| $\boldsymbol{\sigma}_{\boldsymbol{N}}$ | 0.2 | | 2 | |
| $\boldsymbol{\sigma}_{\boldsymbol{R}}$ | 0.16 | | 2 | |
| $\boldsymbol{\sigma}_{\boldsymbol{C}_{\boldsymbol{4}}}$ | 0.14 | | 1.8 | |
| $\boldsymbol{\sigma}_{\boldsymbol{C}_{\boldsymbol{8}}}$ | 0.19 | | 1.8 | |
| $\boldsymbol{\sigma}_{\boldsymbol{V}}$ | 0.5 | | 2.4 | |
| $\boldsymbol{\sigma}_{\boldsymbol{E}}$ | 0.21 | | 9.3 | |
| $\boldsymbol{\sigma}_{\boldsymbol{M}}$ | 0.3 | | 9.6 | |

**Table S7.** Individual parameter estimates for the fits of the model in **eqs. 2-3** in main text (lowest AIC in **Table S5**) to the T cell and virus dynamics. Values of $\bar{\psi}$ for $\beta,\omega_{4},\omega_{8}$ and $I_{50}$ shown here are transformed assuming a blood volume of 3×10^5^ μL (calculated assuming blood:weight ratio of 60mL/Kg and body weight of 5Kg). Shown are individual estimates for animals that continued study after ATI.

|  | **Control** | | | | **WT-Transplant** | | | | **ΔCCR5-Transplant** | | | | | |
| --- | --- | --- | --- | --- | --- | --- | --- | --- | --- | --- | --- | --- | --- | --- |
| **Par.**  **ID** | **Z09087** | **Z09106** | **Z09192** | **Z09204** | **Z09144** | **Z08214** | **A11200** | **Z09196** | **A11219** | **T10187** | **R10159** | **T10173** | **Z11151** | **Z12420** |
| ${\hat{\boldsymbol{r}}}_{\boldsymbol{p}}^{\boldsymbol{j}}$  **(1/day)** | 0.03 | 0.07 | 0.07 | 0.07 | 0.06 | 0.05 | 0.06 | 0.12 | 0.05 | 0.11 | 0.06 | 0.05 | 0.06 | 0.11 |
| ${\hat{\boldsymbol{r}}}_{\boldsymbol{s}}^{\boldsymbol{j}}$  **(1/day)** | 0.06 | 0.07 | 0.09 | 0.06 | 0.08 | 0.12 | 0.07 | 0.15 | 0.10 | 0.10 | 0.08 | 0.06 | 0.06 | 0.07 |
| ${\hat{\boldsymbol{r}}}_{\boldsymbol{m}}^{\boldsymbol{j}}$ **(1/day)** | 0.05 | 0.05 | 0.05 | 0.05 | 0.05 | 0.05 | 0.05 | 0.05 | 0.05 | 0.05 | 0.05 | 0.05 | 0.05 | 0.05 |
| ${\hat{\boldsymbol{r}}}_{\boldsymbol{e}}^{\boldsymbol{j}}$  **(1/day)** | 0.04 | 0.05 | 0.07 | 0.04 | 0.06 | 0.08 | 0.05 | 0.13 | 0.06 | 0.08 | 0.07 | 0.05 | 0.03 | 0.06 |
| ${\hat{\boldsymbol{d}}}_{\boldsymbol{n}}^{\boldsymbol{j}}$  **(1/day)** | 0.01 | 0.02 | 0.02 | 0.01 | 0.01 | 0.01 | 0.01 | 0.02 | 0.02 | 0.02 | 0.03 | 0.02 | 0.01 | 0.02 |
| $\boldsymbol{\lambda}_{\boldsymbol{p}}^{\boldsymbol{j}}$  **(1/day)** | 0.0002 | 0.0002 | 0.0002 | 0.0002 | 0.0002 | 0.0002 | 0.0002 | 0.0002 | 0.0002 | 0.0002 | 0.0002 | 0.0002 | 0.0002 | 0.0002 |
| $\boldsymbol{\lambda}_{\boldsymbol{n}}^{\boldsymbol{j}}$  **(1/day)** | 0.002 | 0.003 | 0.003 | 0.002 | 0.003 | 0.002 | 0.002 | 0.004 | 0.004 | 0.004 | 0.005 | 0.004 | 0.002 | 0.003 |
| $\boldsymbol{\lambda}_{\boldsymbol{s}}^{\boldsymbol{j}}$  **(1/day)** | 0.02 | 0.02 | 0.02 | 0.03 | 0.04 | 0.01 | 0.01 | 0.03 | 0.02 | 0.02 | 0.02 | 0.02 | 0.05 | 0.03 |
| $\boldsymbol{\lambda}_{\boldsymbol{m}}^{\boldsymbol{j}}$ **(1/day)** | 0.08 | 0.08 | 0.08 | 0.08 | 0.08 | 0.08 | 0.08 | 0.08 | 0.08 | 0.08 | 0.08 | 0.08 | 0.08 | 0.08 |
| $\boldsymbol{K}_{\boldsymbol{p}}^{\boldsymbol{j}}$  $\left( \boldsymbol{cells} \right)$ | 10^8.8^ | 10^8.7^ | 10^8.9^ | 10^8.9^ | 10^8.6^ | 10^8.8^ | 10^8.6^ | 10^8.5^ | 10^8.5^ | 10^8.6^ | 10^8.9^ | 10^8.7^ | 10^8.8^ | 10^8.6^ |
| $\boldsymbol{K}_{\boldsymbol{s}}^{\boldsymbol{j}}$  $\left( \frac{\boldsymbol{cells}}{\boldsymbol{\mu L}} \right)$ | 1630.5 | 1356.7 | 2147.8 | 1807.4 | 940.2 | 1488.5 | 1111.4 | 863.4 | 764.5 | 1043.1 | 2036.3 | 1295.9 | 1617.6 | 942.5 |
| $\boldsymbol{K}_{\boldsymbol{m}}^{\boldsymbol{j}}$  $\left( \frac{\boldsymbol{cells}}{\boldsymbol{\mu L}} \right)$ | 999.4 | 1105.0 | 1439.7 | 914.1 | 496.3 | 742.5 | 617.1 | 570.5 | 334.3 | 499.8 | 807.2 | 477.3 | 1094.8 | 523.3 |
| $\boldsymbol{K}_{\boldsymbol{e}}^{\boldsymbol{j}}$  $\left( \frac{\boldsymbol{cells}}{\boldsymbol{\mu L}} \right)$ | 1564.3 | 1301.6 | 2060.6 | 1734.0 | 902.0 | 1428.1 | 1066.3 | 828.3 | 733.4 | 1000.7 | 1953.7 | 1243.3 | 1551.9 | 904.3 |
| $\boldsymbol{k}_{\boldsymbol{T}}^{\boldsymbol{j}}$  **(1/day)** | 0.52 | 0.53 | 0.55 | 0.53 | 0.54 | 0.67 | 0.46 | 0.57 | 0.51 | 0.44 | 0.58 | 0.41 | 0.54 | 0.57 |
| $\boldsymbol{k}_{\boldsymbol{H}}^{\boldsymbol{j}}$  **(1/day)** | 1.65 | 1.65 | 1.65 | 1.65 | 1.61 | 1.71 | 1.63 | 1.69 | 1.63 | 1.70 | 1.83 | 1.70 | 1.59 | 1.63 |
| $\boldsymbol{\beta}^{\boldsymbol{j}}$ $\left( \frac{\boldsymbol{\mu L}}{\boldsymbol{copies*day}} \right)$ | 0.0005 | 0.0002 | 0.0002 | 0.0003 | 0.0001 | 0.0001 | 0.0002 | 0.0002 | 0.0001 | 0.0002 | 0.0001 | 0.0001 | 0.0004 | 0.0003 |
| $\boldsymbol{t}_{\boldsymbol{sa}}^{\boldsymbol{j}}$ **(days)** | 1.1 | 3.5 | 3.1 | 3.8 | 9.7 | 15.6 | 12.9 | 10.5 | 17.3 | 8.3 | 10.1 | 16.5 | 12.7 | 47.3 |
| $\boldsymbol{\pi}^{\boldsymbol{j}}$  **(1/day)** | 10^4.7^ | 10^5.3^ | 10^4.9^ | 10^4.7^ | 10^5.7^ | 10^5.7^ | 10^5.0^ | 10^5.4^ | 10^5.6^ | 10^5.3^ | 10^5.0^ | 10^5.0^ | 10^4.7^ | 10^5.1^ |
| $\boldsymbol{\omega}_{\boldsymbol{4}}^{\boldsymbol{j}}$ $\left( \frac{\boldsymbol{\mu L}}{\boldsymbol{cells*day}} \right)$ | 0.003 | 0.005 | 0.001 | 0.003 | 0.006 | 0.005 | 0.009 | 0.005 | 0.003 | 0.045 | 0.002 | 0.007 | 0.003 | 0.006 |
| $\boldsymbol{\omega}_{\boldsymbol{8}}^{\boldsymbol{j}}$ $\left( \frac{\boldsymbol{\mu L}}{\boldsymbol{cells*day}} \right)$ | 0.00003 | 0.00010 | 0.00008 | 0.00642 | 0.00097 | 0.00183 | 0.00205 | 0.00069 | 0.00050 | 0.00128 | 0.00009 | 0.00108 | 0.00008 | 0.00908 |
| $\boldsymbol{\omega}_{\boldsymbol{8}}^{\boldsymbol{j,ATI}}$ $\left( \frac{\boldsymbol{\mu L}}{\boldsymbol{cells*day}} \right)$ | 0.0001 | 0.0004 | 0.0001 | 0.7414 | 0.0336 | 0.0174 | 1.2605 | 0.0082 | 0.0294 | 0.0007 | 0.0088 | 0.2749 | 0.0081 | 0.4365 |
| $\boldsymbol{d}_{\boldsymbol{h}}^{\boldsymbol{j}}$  **(1/day)** | 0.002 | 0.002 | 0.001 | 0.008 | 0.006 | 0.013 | 0.006 | 0.005 | 0.002 | 0.008 | 0.002 | 0.003 | 0.003 | 0.009 |
| $\boldsymbol{d}_{\boldsymbol{h}}^{\boldsymbol{j,ATI}}$  $\mathbf{(1/day)}$ | 0.005 | 0.012 | 0.004 | 0.257 | 0.457 | 0.753 | 0.384 | 0.150 | 0.179 | 0.027 | 0.019 | 0.045 | 0.182 | 0.043 |
| $\boldsymbol{I}_{\mathbf{50}}^{\boldsymbol{j}}$  $\left( \frac{\boldsymbol{cells}}{\boldsymbol{\mu L}} \right)$ | 5.2 | 11.9 | 65.4 | 0.9 | 1.6 | 7.7 | 1.6 | 5.4 | 13.2 | 0.8 | 27.0 | 5.1 | 15.9 | 0.8 |
| $\boldsymbol{I}_{\mathbf{50}}^{\boldsymbol{j,ATI}}$  $\left( \frac{\boldsymbol{cells}}{\boldsymbol{\mu L}} \right)$ | 18.2 | 34.9 | 264.2 | 3.0 | 5.3 | 29.8 | 5.1 | 17.8 | 45.4 | 3.6 | 93.2 | 19.2 | 62.1 | 2.9 |
